# Constitutive depletion of brain serotonin differentially affects rats’ social and cognitive abilities

**DOI:** 10.1101/2021.09.23.461469

**Authors:** Lucille Alonso, Polina Peeva, Sabrina Stasko, Michael Bader, Natalia Alenina, York Winter, Marion Rivalan

**Affiliations:** Humboldt-Universität zu Berlin, Berlin, Germany; Charité – Universitätsmedizin Berlin, corporate member of Freie Universität Berlin and Humboldt-Universität zu Berlin, Berlin, Germany; Max Delbrück Center for Molecular Medicine in the Helmholtz Association, Berlin, Germany; Institute of Translational Biomedicine, St. Petersburg State University, St. Petersburg, Russia

**Author notes:** Corresponding authors: Marion Rivalan, E-mail address, Postal address: Charité – Universitätsmedizin Berlin, Exzellenzcluster NeuroCure, Animal Behaviour Phenotyping Facility, CharitéCrossOver, Virchowweg 6, 10117 Berlin, Germany Natalia Alenina, E-mail address, Postal address: Max Delbrück Centrum für Molekulare Medizin (MDC), Robert-Rössle-Straße 10, 13125 Berlin, Germany.

**Keywords:** Serotonin, TPH2 knockout, social interaction, decision-making, automated home-cage, rat behavioral profile

## Abstract

**Background:** Central serotonin is an essential neuromodulator for mental disorders. It appears a promising transdiagnostic marker of distinct psychiatric disorders and a common modulator of some of their key behavioral symptoms. We aimed to identify the behavioral markers of serotonergic function in rats and compare them to human deficits.

**Methods:** We applied a comprehensive profiling approach in adult male *Tph2*^*−/−*^ rats constitutively lacking central serotonin. Under classical and ethological testing conditions, we tested each individual’s cognitive, social and non-social abilities and characterized the group organization (i.e. social network, hierarchy). Using unsupervised machine learning, we identified the functions most dependent on central serotonin.

**Results:** In classical procedures, *Tph2*^*−/−*^ rats presented an unexpected normal cognitive profile. Under the complex and experimenter-free conditions of their home-cage, the same *Tph2*^*−/−*^ rats presented drastic changes in their daily life. Brain serotonin depletion induced compulsive aggression and sexual behavior, hyperactive and hypervigilant stereotyped behavior, reduced self-care and body weight, and exacerbated corticosterone levels. Group-housed *Tph2*^*−/−*^ rats showed strong social disorganization with disrupted social networks and hierarchical structure, which may arise from communication deficits and cognitive blunting.

**Conclusions:** Serotonin depletion induced a profile reminiscent of the symptomatology of impulse control and anxiety disorders. Serotonin was necessary for behavioral adaptation to dynamic social environments. In classical testing conditions, our animal model challenged the concept of an essential role of serotonin in decision-making, flexibility, and impulsivity, although developmental compensations may have occurred. These contrasting findings highlight the need to generalize the evaluation of animal models’ multidimensional functions within the complexity of the social living environment.

## Introduction

The complex nature of psychiatric disorders makes them some of the least understood and most incapacitating of all pathological conditions (1–3). A challenge for biomedical research today is to develop efficient and specific treatments that can reverse dysfunctional conditions and improve psychiatric patients’ quality of life. However, the current diagnosis of mental disorders lacks biological markers specific to given pathological conditions (2). Beyond the categorical classification of psychiatric disorders, the search for combinations of behavioral symptoms associated with a specific biological profile is necessary for identifying neurocognitive markers of mental disorders (4–6).

The monoamine serotonin (5-hydroxytryptamine) is a neuromodulator of the central nervous system (CNS). In the CNS, its synthesis is restricted to the raphe nuclei neurons, which innervate the whole brain with a vast axonal network (7–10). Serotonin, through its action on numerous post- and presynaptic receptors (11), is essential for mood regulation and treating mood disorders (anxiety, bipolar, and depressive disorders; 9,12) and other neuropsychiatric disorders, such as addiction (13–15), attention deficit hyperactivity disorder (16), suicidal behavior (17,18), obsessive-compulsive disorder (19,20), psychopathy (21), and other aggression-related disorders (22,23). At the behavioral level, serotonin is known to be critical in modulating several executive functions and aspects of social behavior. Disadvantageous decisions (24,25), impulsive choices and actions (26–28), inflexibility (26,27,29), aggression, and socially inappropriate behavior (30,31) are characteristic impairments of affective, impulse control, or substance-related disorders (32–39). Similarly, such cognitive and social deficits are induced in non-clinical humans and rodents after experimental reduction of serotonin levels (40–47).

Overall, the serotonergic system appears a promising transdiagnostic marker of apparently distinct psychiatric disorders and a common modulator of some of their key behavioral symptoms. Despite the appeal to reduce mental disorders to impairments studied in isolation, the reality is that the complexity of human mental disorders cannot be explained only in terms of their components, as their interaction plays a critical role in the emergence of the pathology (48–51). Using a multidimensional profiling approach (52), we studied the effect of brain serotonin depletion on the expression of several cognitive, social, and affective functions in the same individual. We aimed to expose the characteristic profile associated with serotonin dysfunction, identify which functions were most affected by the absence of brain serotonin, and compare this profile to mental conditions observed in humans.

Genetic modifications are among the most specific methods to target central serotonin in animals. In the recently created line of tryptophan hydroxylase 2 knockout rats (*Tph2*^*−/−*^; 53), the constitutive absence of the TPH2 enzyme disables the production of brain serotonin (54). *Tph2*^*−/−*^ pups display delayed growth and impaired autonomic responses (53), which normalize at adult age. At the behavioral level, *Tph2*^*−/−*^ rats showed increased aggression in the resident intruder paradigm (55). However, more subtle social and cognitive deficits remain to be characterized.

Based on previous studies where executive and social functions were individually tested after pharmacological, genetic, or dietary alteration of central serotonin, we hypothesized that the absence of serotonin would simultaneously alter the rats’ cognitive and executive functions, social abilities, activity level, and affective responses in both classical testing contexts and more dynamic home-cage environments. We used the ethological environment of a version the visible burrow system (VBS; 52) to identify novel real-life markers of serotonergic function (56). The *Tph2*^*−/−*^ phenotype was characterized by multiple behavioral changes only detected in the dynamic social context. With unsupervised machine learning we uncovered that the most critical impairments in these animals resembled transdiagnostic symptoms of impulse control disorders.

## Material and methods

### Animals

We used 48 *Tph2*^*+/+*^ and 30 *Tph2*^*−/−*^ male rats [TPH2-ZFN; (53); Table S1] housed in pairs of the same genotype in two temperature-controlled rooms (22°C–24°C and 45%–55% humidity) with inverted 12-hour light-dark cycles. Rats were individually marked subcutaneously with radio-frequency identification (RFID) tags. They had *ad libitum* access to water. They were maintained at 95% of their free-feeding weight during operant training and testing and otherwise fed *ad libitum*. Rats were between 8 and 14 weeks old when first trained in the operant procedures. Descriptions of handling and housing methods are provided in the supplemental methods section. The study was reported in accordance with the ARRIVE Guidelines [ARRIVE Checklist, (57)].

### Ethics

All procedures followed the national regulations in accordance with the European Union Directive 2010/63/EU. The protocols were approved by the local animal care and use committee and under the supervision of the animal welfare officer of our institution.

### Behavioral tests

Due to space limitations, details of apparatus, methods, and parameters of this study are provided in the supplemental methods section. Unless stated otherwise, rats were trained and tested following established procedures described previously (52). The order of tests is shown in Fig. 1.

**Figure 1.**
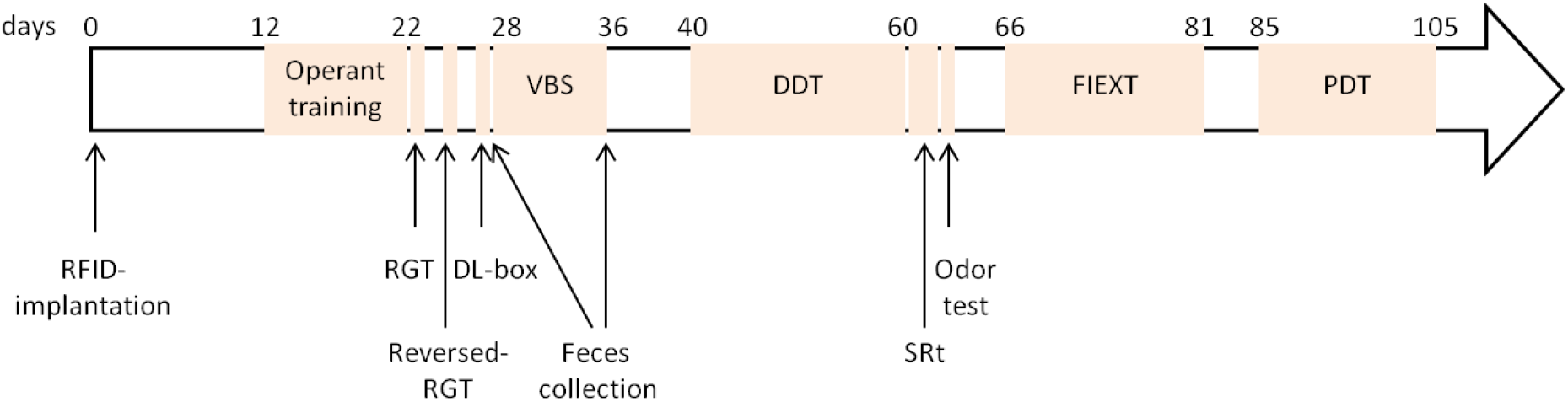
Order of tests. We used 8 *Tph2*^*+/+*^ and 5 *Tph2*^*−/−*^ cohorts, six animals each. Radio-frequency identification (RFID), rat gambling task (RGT), dark-light box test (DL-box), automated visible burrow system (VBS), delay discounting task (DDT), social recognition test (SRt), odor discrimination test (odor test), fixed-interval and extinction schedule of reinforcement test (FIEXT), probability discounting task (PDT).

### Operant system

Four operant cages were used (Imetronic, France) with either a curved wall equipped with one to four nose-poke holes or a straight wall equipped with one central lever, depending on the test. On the opposite wall was a food magazine connected to an outside pellet dispenser filled with 45 mg sweet pellets (5TUL, TestDiet, USA).

### Complex decision-making in the rat gambling task (RGT)

In the RGT, two nose-poke holes on one side of the wall were rewarded with a large reward (two pellets) and associated with unpredictable, long timeouts (222 and 444 s with probabilities of occurrence of 1/2 and 1/4, respectively); within the 60 min of testing, these options were disadvantageous. Two nose-poke holes on the other side of the same wall were rewarded with one pellet and associated with unpredictable, short timeouts (6 and 12 s with probabilities of occurrence 1/2 and 1/4, respectively); these options were advantageous. The mean latency to visit the feeder after a choice and percentages of advantageous choices per 10 min and for the last 20 min of the test were recorded.

### Cognitive flexibility in the reversed rat gambling task (reversed-RGT)

In the reversed-RGT, the two disadvantageous options were spatially switched with the two advantageous options of the RGT, and the flexibility score was recorded.

### Cognitive impulsivity in the delay discounting task (DDT)

In the DDT, one nose-poke hole (NP1) was associated with a small immediate reward (1 pellet) and a second nose-poke hole (NP5; 25 cm between the two holes) with a large (5 pellets) delayed (0, 10, 20, 30, or 40 s) reward. The preference for the large reward at each delay was recorded.

### Risky decision-making in the probability discounting task (PDT)

In the PDT, one hole (NP1) was associated with a small and sure reward (1 pellet), and a second nose-poke hole (NP5) was associated with a large (5 pellets) and uncertain (P = 1, 0.66, 0.33, 0.20, 0.14, or 0.09) reward. The preference for the large reward at each probability was recorded.

### Social recognition task (SRt)

The SRt occurred in a square open arena. In one corner of the arena was a smaller cage where an unfamiliar rat could be placed that the test rat could smell through a grid wall. The test rat could first explore the empty arena for 15 min of habituation. An unfamiliar conspecific was then placed in the cage, and the test rat was allowed to explore for three consecutive trials of 5 min (E1, E2, E3) with a 10-min break between encounters. The durations of interaction with the empty cage (habituation) and the cage with the conspecific were recorded.

### Spontaneous behaviors, space occupation, hierarchy, and social network analysis in the automated VBS

The VBS comprised an open home-cage arena where food and water were available at all times and a burrow system connected to the open area by two tunnels. The burrow system was kept in the dark throughout the test (infrared-transparent black plastic) and comprised a large and a small chamber connected by tunnels. A grid of 32 RFID detectors (PhenoSys, Germany) was placed underneath the VBS to automatically determine individual animal positions. Six rats of the same genotype were housed together in the VBS. The duration of the VBS housing was reduced from 7 to 4 days for the *Tph2*^*−/−*^ animals after the first group to limit differences in weight loss between groups (58). The behaviors of the first 4 h of each dark and light phase were scored by trained experimenters using videos (Table 1). All aggressive behaviors except “struggling at feeder” were grouped under “general aggression” (Table 1). Individual rats’ activity (distance traveled), place preference, time spent in the open area, and body weight change were recorded.

**Table 1.**
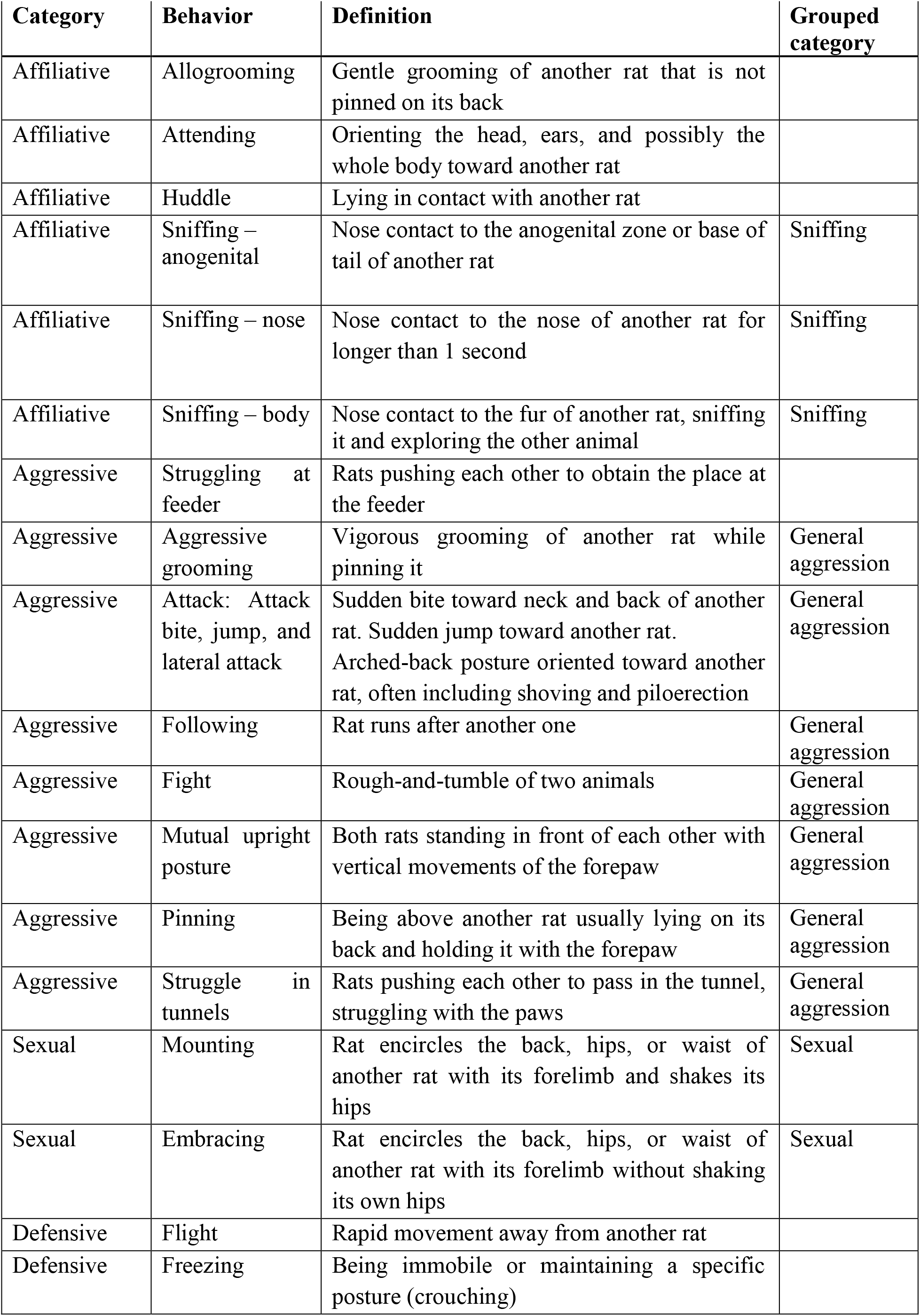
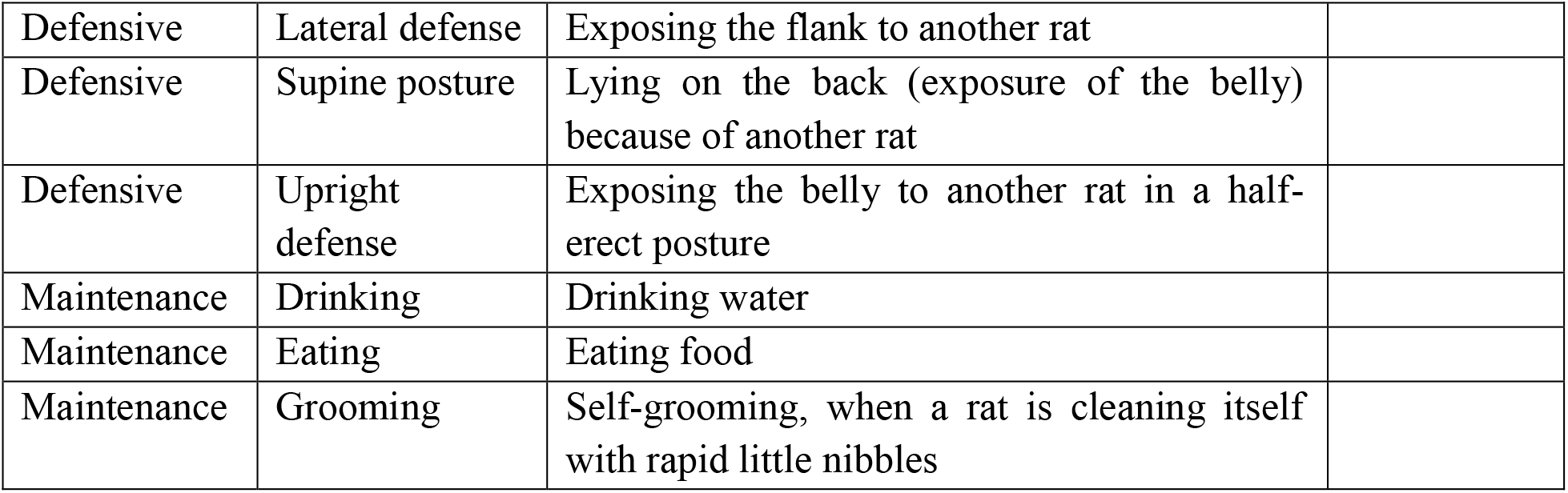
Ethogram of the behaviors scored in the VBS [based on (59–62)].

| **Category** | **Behavior** | **Definition** | **Grouped category** |
| --- | --- | --- | --- |
| Affiliative | Allogrooming | Gentle grooming of another rat that is not pinned on its back |  |
| Affiliative | Attending | Orienting the head, ears, and possibly the whole body toward another rat |  |
| Affiliative | Huddle | Lying in contact with another rat |  |
| Affiliative | Sniffing – anogenital | Nose contact to the anogenital zone or base of tail of another rat | Sniffing |
| Affiliative | Sniffing – nose | Nose contact to the nose of another rat for longer than 1 second | Sniffing |
| Affiliative | Sniffing – body | Nose contact to the fur of another rat, sniffing it and exploring the other animal | Sniffing |
| Aggressive | Struggling at feeder | Rats pushing each other to obtain the place at the feeder |  |
| Aggressive | Aggressive grooming | Vigorous grooming of another rat while pinning it | General aggression |
| Aggressive | Attack: Attack bite, jump, and lateral attack | Sudden bite toward neck and back of another rat. Sudden jump toward another rat. Arched-back posture oriented toward another rat, often including shoving and piloerection | General aggression |
| Aggressive | Following | Rat runs after another one | General aggression |
| Aggressive | Fight | Rough-and-tumble of two animals | General aggression |
| Aggressive | Mutual upright posture | Both rats standing in front of each other with vertical movements of the forepaw | General aggression |
| Aggressive | Pinning | Being above another rat usually lying on its back and holding it with the forepaw | General aggression |
| Aggressive | Struggle in tunnels | Rats pushing each other to pass in the tunnel, struggling with the paws | General aggression |
| Sexual | Mounting | Rat encircles the back, hips, or waist of another rat with its forelimb and shakes its hips | Sexual |
| Sexual | Embracing | Rat encircles the back, hips, or waist of another rat with its forelimb without shaking its own hips | Sexual |
| Defensive | Flight | Rapid movement away from another rat |  |
| Defensive | Freezing | Being immobile or maintaining a specific posture (crouching) |  |
| Defensive | Lateral defense | Exposing the flank to another rat |  |
| Defensive | Supine posture | Lying on the back (exposure of the belly) because of another rat |  |
| Defensive | Upright defense | Exposing the belly to another rat in a half-erect posture |  |
| Maintenance | Drinking | Drinking water |  |
| Maintenance | Eating | Eating food |  |
| Maintenance | Grooming | Self-grooming, when a rat is cleaning itself with rapid little nibbles |  |

### Glicko rating

For each VBS group, the social ranking of the rats was defined using a Glicko rating system (63,64). Briefly, the individual rank was dynamically updated for each individual following the outcome of each aggressive and sexual interaction within the group. The normalized number of change points and maximum rating contrast were recorded.

### Roaming entropy

The roaming entropy (RE) within the VBS is the probability that an individual will be at a certain place at a given time. The RE calculation was based on the method described by Freund et al. (65). In the VBS, the spatial dispersion of the rats was evaluated through the total and daily RE.

### Social network analysis

We used social network analysis methods to expose qualitative aspects of the social interactions in the VBS, such as information transmission or power distribution within each group. For each behavioral network (huddling, sniffing, struggling at the feeder, general aggression, and sexual behaviors), global parameters (density, average path length, out-degree centralization) and individual parameters (e.g. hub centrality) were calculated.

### Feces collection and corticosterone metabolite measurements

Corticosterone metabolite concentrations were measured before and after VBS housing.

### Statistics

R [version R – 3.6.1; (66)] and R studio (version 1.1.456) were used for statistical analyses. All data and scripts are available in the Zenodo repository: https://doi.org/10.5281/zenodo.4912528. Briefly, we used the Wilcoxon rank sum test for genotype comparisons, Fisher’s exact test for proportion comparisons, and the one-sample t-test and Wilcoxon rank sum test for comparison with a theoretical value. Linear mixed models (lmer) and generalized linear models with Markov chains (glmmMCMC) were used for repeated measures analysis. Random forest (RF) and principal component analysis (PCA) were used to identify the functions most affected by brain serotonin depletion in tests. One *Tph2*^*+/+*^ animal was excluded from the RGT because it did not sample the options. One *Tph2*^*−/−*^ animal was excluded from the odor discrimination test because it did not explore the open field. One group of six *Tph2*^*+/+*^ rats was excluded from RE analysis due to a grid malfunction on days 1 and 2. Due to space limitations, statistics are provided in a supplemental table where more than two results must be reported.

## Results

### Central serotonin deficiency does not affect decision-making, cognitive flexibility, sensitivity to reward, motor impulsivity, social memory, and anxiety

All the animals started the RGT without preference for either option (first 10 min, Fig. 2A) and preferentially chose the advantageous options over the disadvantageous ones after 10 minutes of the test (Fig. 2A, one-sample t-test, 20 min: +/+: 0.95CI [53.7, 73.9], p-value = 0.008; −/−: 0.95CI [59.5, 85.2], p-value < 0.001 and Table S2). In both *Tph2*^*+/+*^ and *Tph2*^*−/−*^ groups, this dynamic was driven by a majority of good decision-makers (GDMs; Fig. 2B and Fig. S1). Unexpectedly, both groups presented the same proportion of good (+/+: 74%; −/−: 73%), intermediate (+/+: 9%; −/−: 10%), and poor decision-makers (PDMs, +/+: 17%; −/−: 17%; Fig. 2B). Regardless of their genotype but consistent with their typical decision-makers’ profile (67), PDMs were faster to collect rewards after a choice compared to GDMs (Fig. 2C, Wilcoxon rank sum test, *PDMs vs. GDMs*: *+/+*: W = 203, p-value = 0.049, −/−: W = 89, p-value = 0.033). PDMs were incapable of flexibility in the reversed-RGT test (Fig. 2D; Wilcoxon rank sum test, *PDMs vs. GDMs*: *+/+*: W = 217, p-value = 0.016, −/−: W = 90.5, p-value = 0.028). *Tph2*^*+/+*^ and *Tph2*^*−/−*^ GDMs made either flexible choices (40% and 45%, respectively), inflexible choices (40% and 45%), or were undecided (20% and 10%, Fig. 2D). GDMs and PDMs did not differ in any other tests or between genotypes (Table S3). For the remainder of the study, only genotype comparisons are presented. In the DDT (Fig. 2E) and PDT (Fig. 2F), rats’ preference for the large reward progressively decreased as the associated discounting factor (delay or uncertainty) increased. Rats of both genotypes switched preference for the (immediate) smaller reward at delay 20 s [Fig. 2E, lmer, *delay*: F(4, 289) = 1, p-value < 0.001] and at probability 20% [Fig. 2F, lmer, *probability*: F(5, 230) = 193, p-value < 0.001]. In the DDT, *Tph2*^*−/−*^ rats presented a smaller total area under the curve (AUC) than *Tph2*^*+/+*^ animals (Fig. 2E inset, Wilcoxon rank sum test, W = 916, p-value = 0.044). In the PDT, both genotypes presented similar AUC (Fig. 2F inset, Wilcoxon rank sum test, W = 372, p-value = 0.085). Animals of both genotypes presented similar anticipatory and perseverative responses during the fixed-interval and extinction phases of the FIEXT schedule of test (Fig. S2). Despite a similar social preference for an unfamiliar partner (E1, Fig. 2G, and Fig. S3) and recognition abilities (E2, E3, Fig. 2G, and Fig. S3) in both groups, *Tph2*^*−/−*^ rats presented a higher interest in the social partner than the *Tph2*^*+/+*^ rats [Fig. 2G, lmer, *genotype*: F(1, 40) = 8, p-value = 0.006]. *Tph2*^*−/−*^ and *Tph2*^*+/+*^ rats showed similar abilities in the odor discrimination test (Fig. S4). Anxiety and risk-taking levels in the dark-light box were similar between genotypes, although *Tph2*^*−/−*^ rats showed high variability in responses (Fig. 2H and I).

**Figure 2.**
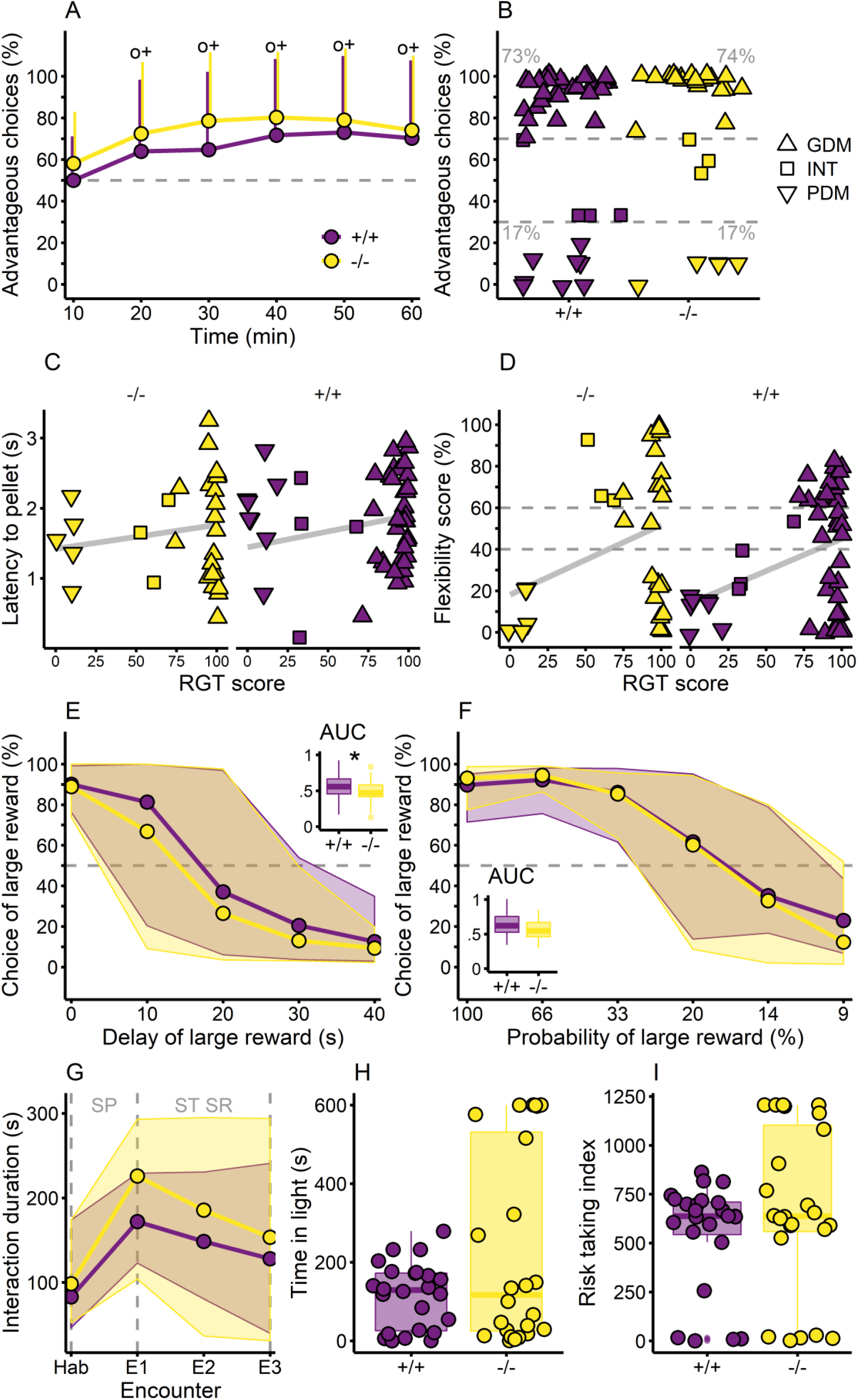
Cognitive abilities of the *Tph2*^*+/+*^ and *Tph2*^*−/−*^ rats. **A**. Advantageous choices in the rat gambling task (RGT). Lines indicate mean + SD, one-sample t-test compared to 50% with ° p-value < 0.05 for +/+ and ^+^ p-value < 0.05 for −/−. **B**. Individual (mean) scores during the last 20 min of the RGT. The dashed lines at 70% and 30% of advantageous choices visually separate good decision-makers (GDMs, above 70% of advantageous choices in the last 20 min, upward triangle), intermediates (INTs, between 30% and 70% of advantageous choices in the last 20 min, square), and poor decision-makers (PDMs, below 30% of advantageous choices in the last 20 min, downward triangle). **C**. Latency to collect the reward in the RGT after a choice for GDMs (upward triangle), INTs (square), and PDMs (downward triangle). Linear regression (grey line) representing the positive correlation. **D**. Flexibility scores in the reversed-RGT corresponding to the preference for the new location of the preferred option in the RGT for GDMs (upward triangle), INTs (square), and PDMs (downward triangle). Linear regression (grey line) representing the positive correlation. The dashed lines at 60% and 40% visually separate flexible individuals (above 60%) from inflexible individuals (below 40%). The flexibility score is the preference for the location of the non-preferred option during the RGT. **E**. Choice of the large reward option as a function of the delay in reward delivery in the DDT. Lines show medians, and shaded areas show 5^th^ to 95^th^ percentiles. The dashed line indicates the 50% chance level. Inset showing the area under the curve (AUC) for the preference for the large reward, Wilcoxon rank sum test between +/+ and −/−, ^*^ p-value < 0.05. **F**. Choice of the large reward option as a function of the probability of reward delivery in the PDT. Lines show medians, and shaded areas show 5^th^ to 95^th^ percentiles. Dashed line shows 50% chance level. Inset showing the AUC for the preference for the large reward. **G**. Duration of interaction in the SRt. Lines show the medians, and shaded areas show the 5^th^ to 95^th^ percentiles, social preference (SP), short-term social recognition (ST SR), habituation with empty cage (Hab), successive encounters with same conspecific placed in the small cage (E1–3). **H**. Time in the open part of the DL-box. Individual data over the boxplot. **I**. Risk-taking index for the DL-box test. Individual data over the boxplot. Panels A–D: +/+ n = 47, −/− n = 30, E: +/+ n = 48, −/− n = 30, F: +/+ n = 24, −/− n = 24, G: +/+ n = 30, −/− n = 30, H–I: +/+ n = 24, −/− n = 24. *Tph2*^*+/+*^ in purple and *Tph2*^*−/−*^ in yellow.

### Central serotonin deficiency disrupts daily activity, place preference, body weight, and corticosterone levels of group-housed rats within the VSB

In the VBS, *Tph2*^*−/−*^ rats were more active than *Tph2*^*+/+*^ rats in reaction to novelty (Fig. 3A, post-hoc test after lmer, day 1 – dark phase: Standard error = 20, z-value = 7, p-value < 0.001) and over days (Fig. 3A, glmmMCMC, *genotype*: post mean = 8.32, credible interval [5.97, 11.03], p-value < 0.001). Circadian fluctuation of day/night activity was preserved in both groups (Fig. 3A, glmmMCMC, *phase*: post mean = -9.14, credible interval [-9.95, -8.42], p-value < 0.001) although it was less pronounced for *Tph2*^*−/−*^ during light phases (glmmMCMC, *genotype x phase*: post mean = 4.13, credible interval [2.83, 5.81], pMCMC < 0.001). According to their RE index, the *Tph2*^*−/−*^ rats used cage space less evenly than the *Tph2*^*+/+*^ rats overall (Fig. 3B, Wilcoxon rank sum test, W = 1183, p-value < 0.001) and over days [Fig. 3C, lmer, *genotype*: F(1, 19) = 27, p-value < 0.001]. They were detected less often at the feeding and drinking areas and in the large chamber than the *Tph2*^*+/+*^ rats (Fig. 3D, purple zones on heatmaps). They stayed more in the covered tunnels close to the open area (burrow area) and in the center of the open area (Fig. 3D, yellow areas). *Tph2*^*−/−*^ rats lost more weight during the VBS stay than *Tph2*^*+/+*^ rats (Fig. 3E, Wilcoxon rank sum test, W = 1397, p-value < 0.001). Only in *Tph2*^*−/−*^ rats, VBS housing largely increased the corticosterone metabolite level [Fig. 3F, lmer, *interaction genotype x time*: F(1, 94) = 69, p < 0.001].

**Figure 3.**
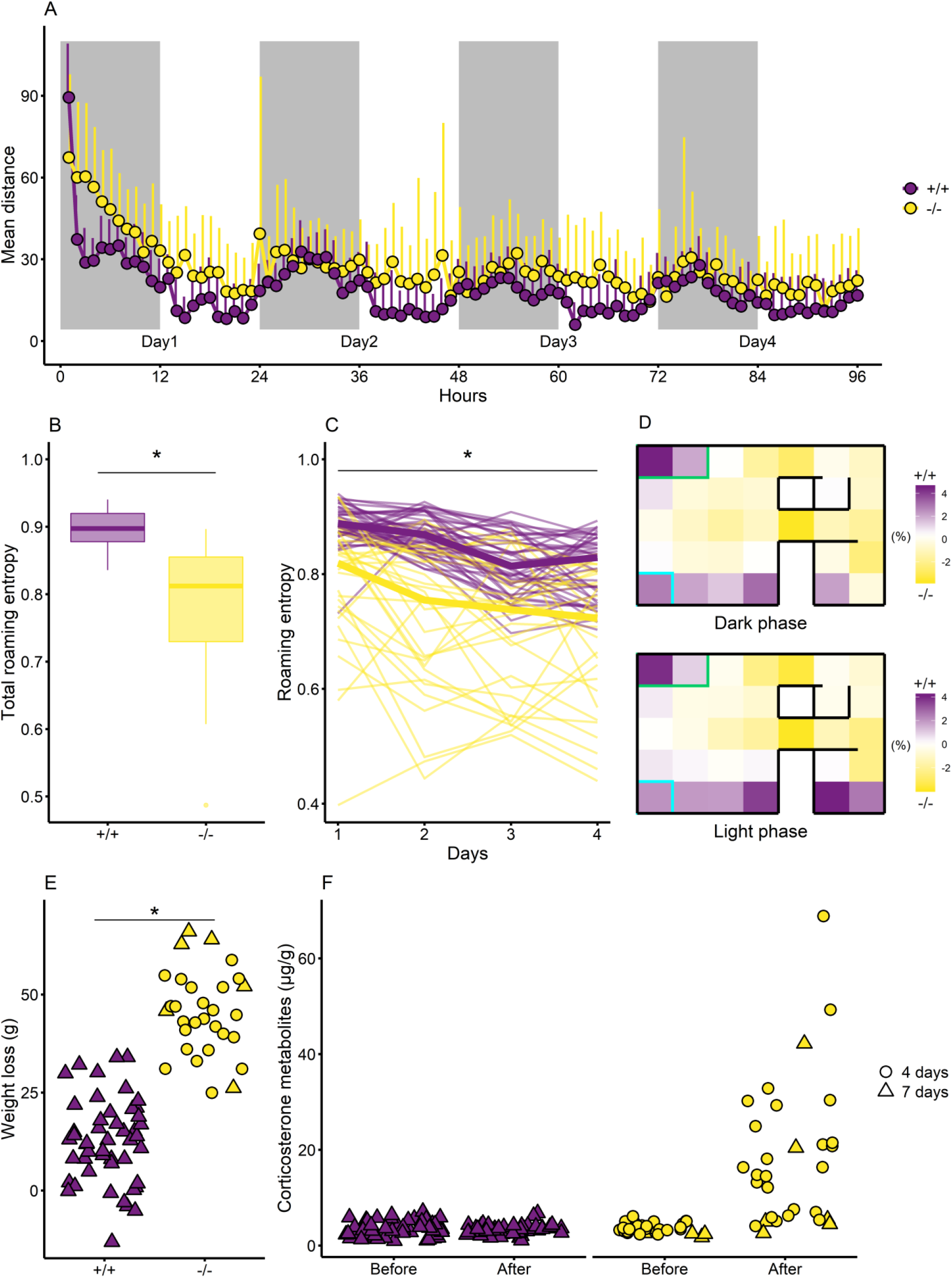
Activity, roaming entropy, and place preference of the *Tph2*^*+/+*^ and *Tph2*^*−/−*^ rats in the automated visible burrow system (VBS). **A**. Activity as mean index of distance traveled in arbitrary unit per hour. Lines indicate mean + SD. **B**. Total roaming entropy, Wilcoxon rank sum test between +/+ and −/−, ^*^p-value < 0.05. **C**. Roaming entropy per day, thick curves indicate the median values, and thin curves indicate the individual values, lmer between +/+ and −/−, ^*^p-value < 0.05. **D**. Difference in place preference (frequency of detections) between +/+ and −/− over 4 days of VBS housing. Purple indicates a higher preference of the +/+ and yellow indicates a higher preference of the −/− for each of the 32 zones of the VBS (corresponding to the 32 RFID detectors located beneath the VBS cage). Rectangles indicate the locations of the feeder (green) and water bottle (cyan). **E**. Weight loss in grams after the stay in the automated VBS. A 4-day stay is indicated with circles, and a 7-day stay is indicated with triangles, Wilcoxon rank sum test between +/+ and −/−, ^*^p-value < 0.05. **F**. Corticosterone metabolites in µg/g of feces before and after VBS housing for both genotypes. A 4-day stay is indicated with circles, and a 7-day stay is indicated with triangles; post-hoc test after lmer between before −/− and after −/− (Standard error = 1.4, z-value = 10.5, p-value < 0.001), ^*^p-value < 0.05. Panels A and D–F: +/+ n = 48, −/− n = 30 and B–C: +/+ n = 42, −/− n = 30. *Tph2*^*+/+*^ in purple and *Tph2*^*−/−*^ in yellow.

### Central serotonin deficiency disrupts social behaviors, social networks, group organization, and hierarchy in the VBS

Overall, *Tph2*^*−/−*^ animals showed less huddling, eating, struggling at the feeder, and grooming behaviors than *Tph2*^*+/+*^ animals and more general aggression, exploratory (sniffing), and sexual behaviors (Fig. 4A, Wilcoxon rank sum test in Table S4 and all behaviors are presented in Fig. S5). On day 1, for aggression and sexual behavior, *Tph2*^*−/−*^ networks were more dense, with most pairs of rats displaying these behaviors, while fewer pairs connected for huddling and struggling at the feeder (Fig. 4B, lmer in Table S5) compared to controls. On the following days and by day 4, the *Tph2*^*−/−*^ network densities for huddling (Fig. 4C-left), sniffing, and general aggression (Fig. 4C-right) normalized to the level of the *Tph2*^*+/+*^ networks; network densities for sexual behaviors always remained higher for *Tph2*^*−/−*^ and for struggling at the feeder remained stable for both genotypes (Fig. 4B; lmer in Table S5). The average path length (mean number of steps between any pair of the network) indicated similar results to density, and the out-degree centralization (distribution of out-interactions) was low for all networks (median at 0.20, Fig. S6). In both genotypes, individual hierarchical ranks emerged progressively (Fig. 4D). The rats’ final Glicko ratings were broadly distributed below and above the initial rating score (Fig. 4D), with one dominant individual identified in each group (except for one *Tph2*^*−/−*^ group with two dominant individuals, Fig. S7). The two hierarchical scales, the non-aggression Blanchard dominance and the Glicko rating scores, correlated positively in *Tph2*^*+/+*^ (r = 0.30, p-value = 0.0405) and negatively in *Tph2*^*−/−*^ (r = -0.45, p-value = 0.0132). Compared to *Tph2*^*+/+*^, *Tph2*^*−/−*^ dominant animals were more aggressive toward subordinates (higher rank divergence; Fig. 4E, Wilcoxon rank sum test, W = 0, p-value = 0.0015) and the *Tph2*^*−/−*^ group’s hierarchy was more unstable (higher number of change points; Fig. 4F, Wilcoxon rank sum test, W = 453, p-value = 0.0061). Finally, in *Tph2*^*+/+*^ rats, the higher the Glicko rating, the higher the hub centrality in the general aggression network (r = 0.40, p-value = 0.0051). This correlation was not found in *Tph2*^*−/−*^ rats (r = 0.04, p-value = 0.8543), indicating that the dominant’s aggression did not influence this network.

**Figure 4.**
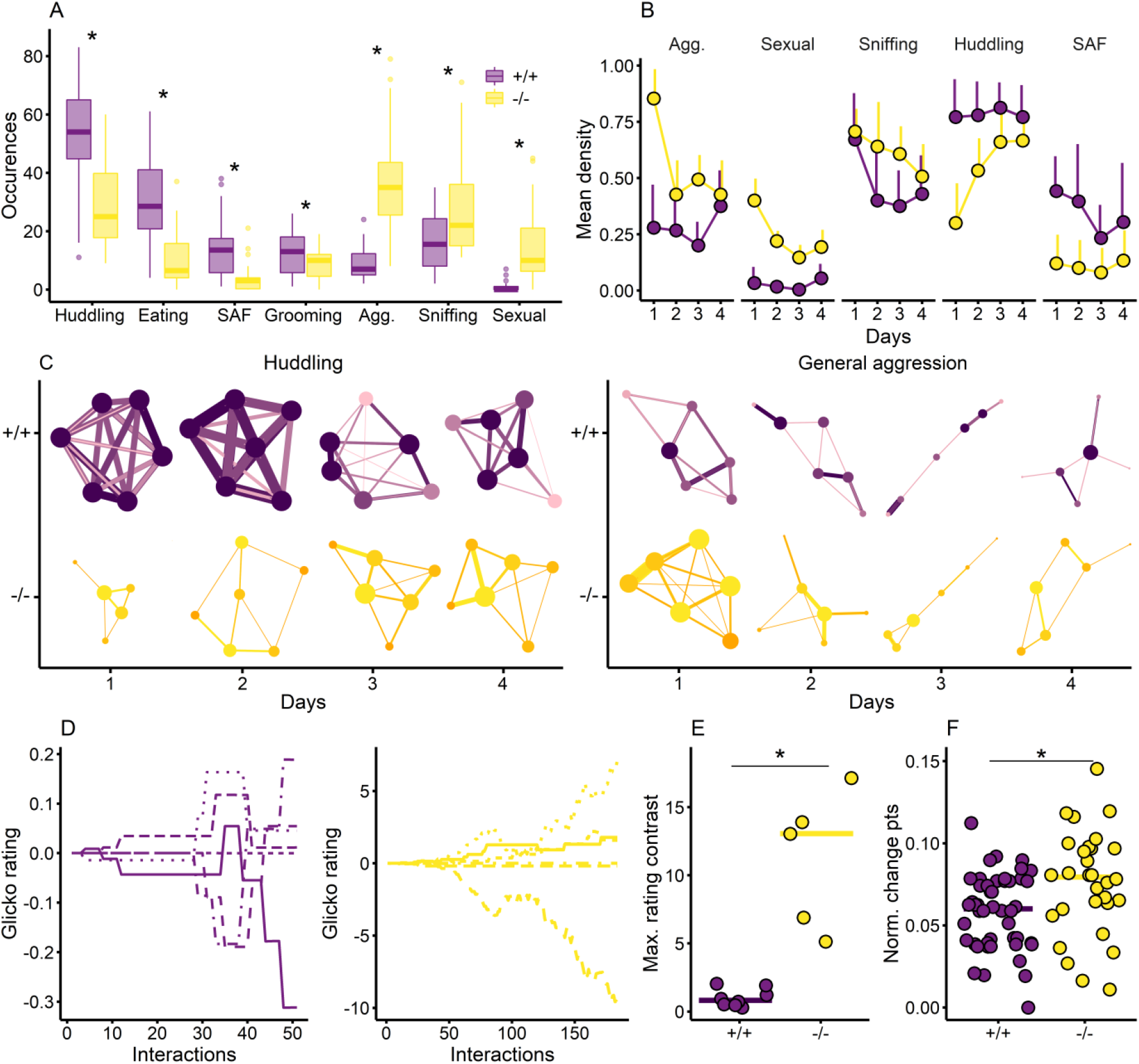
Social abilities and dominance of *Tph2*^*+/+*^ and *Tph2*^*−/−*^ in the automated VBS. **A**. Number of occurrences of behaviors in 4 days in the VBS for the most expressed behaviors, struggling at the feeder (SAF), general aggression including all aggressive behaviors except struggling at the feeder (Agg.), all sniffing behaviors (Sniffing), sexual behaviors including embracing and mounting behaviors (Sexual; Table 1). Wilcoxon rank sum test between *Tph2*^*+/+*^ and *Tph2*^*−/−*, *^p-value < 0.05. **B**. Social network density along days. Lines indicate mean + SD. The network density is the proportion of potential connections in the network that are existing connections between rats; the development of the social network density over days can be visualized by viewing the number of edges in the networks in the panel. **C**. Social network representation from days 1 to 4 for huddling (left) and aggression (right) behaviors. The color intensity and thickness of the edges represent the number of behaviors exchanged (weight), and the color intensity and size of the nodes represent the number of edges received and sent out (node-degree). In the huddling networks, in *Tph2*^*+/+*^, the density was the highest at day 1 and remained high over days as shown by the number of edges and large node size; in *Tph2*^*−/−*^, the density was the lowest at day 1 and increased over days. In the aggression networks, in *Tph2*^*+/+*^, the density was stable and low over days; in the *Tph2*^*−/−*^ group, the density of connection strongly decreased after day 1. **D**. Glicko rating representation for the six individuals of one *Tph2*^*+/+*^ group (left) and for the six individuals of one *Tph2*^*−/−*^ group (right). **E**. Maximum difference in the final Glicko rating between the lowest and highest individuals (Max. rating contrast) for each group, Wilcoxon rank sum test between *Tph2*^*+/+*^ and *Tph2*^*−/−*, *^p-value < 0.05. **F**. Individual proportion of Glicko rating change points, normalized number of change points to the total number of interaction (Norm. change pts); a change point indicates an increase or decrease in the individual rating, Wilcoxon rank sum test between *Tph*2^*+/+*^ and *Tph2*^*−/−*, *^p-value < 0.05. Panels A, B, and F: +/+ n = 48, −/− n = 30, panel E: +/+ n = 8 groups, −/− n = 5 groups and panels C and D representative groups of each genotype. *Tph2*^*+/+*^ in purple and *Tph2*^*−/−*^ in yellow.

### Central serotonin deficiency differentially impacts cognitive abilities and group-housed behaviors

Among all measured behaviors, those most impacted by the lack of brain serotonin were identified using a RF classifier (with an average accuracy of 98.5%, SD = 0.54, Table S6) and confirmed by a PCA (Table S7). The PCA revealed a clear separation of the genotypes along its first dimension (Fig. 5A-left). The variables contributing the most to dimension 1 were also those discriminating the best between genotypes using the RF classifier (Fig. 5B, Table S8 and S9). Dimension 1 was mainly loaded by weight loss, maintenance (drinking, eating, grooming) behavior, RE, corticosterone variation, and defensive and sexual behaviors (Fig. 5A-right). From the RF, the other relevant variables comprised total distance traveled, Glicko rating score, affiliative (allogrooming, attending, huddling, sniffing) and aggressive (struggling at the feeder and general aggression; Table 1) behaviors, and the presence in the VBS open area (Fig. 5B). None of the cognitive variables predicted the animals’ genotypes (Fig. 5A-right and B).

**Figure 5.**
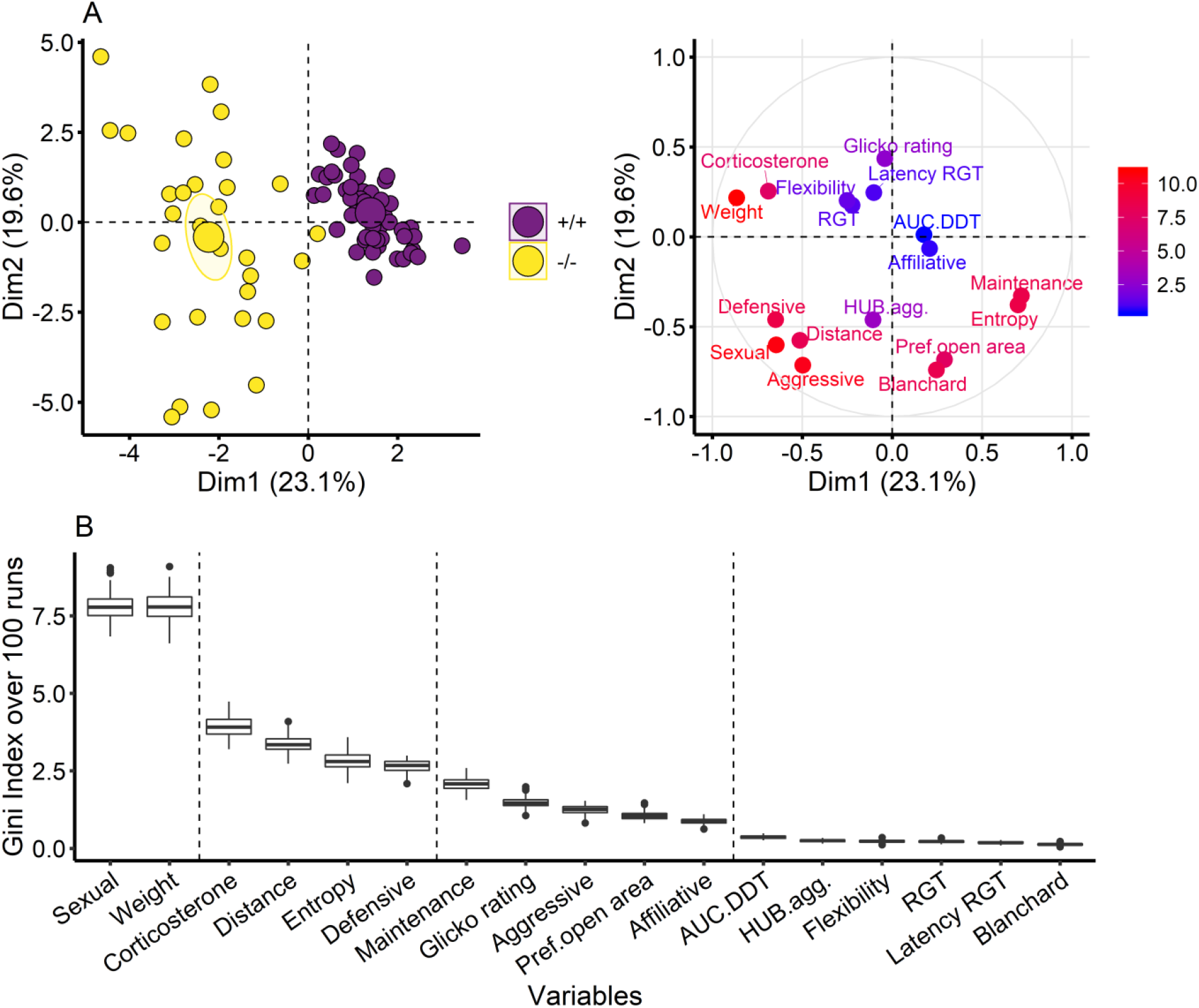
Principal component analysis (PCA) and random forest (RF) classification. **A-left**. Separation of the genotypes along dimension 1 but not along dimension 2 of the principal component analysis, *Tph2*^*+/+*^ in purple and *Tph2*^*−/−*^ in yellow; large symbols show group centroids and ellipses show the 0.95 confidence interval **A-right**. Contribution of the variables to dimensions 1 and 2 of the principal component analysis; higher contribution with warmer color (red) and lower contribution with colder color (blue). **B**. Gini index of the RF classification over 100 runs indicating the importance of the variable = the genotype dissimilarity. The dashed line indicates the groups of variables resulting from the k-means clustering of the Gini indexes over 100 runs. Total occurrences of sexual behaviors (Sexual), percentage of weight variation (Weight), percentage of corticosterone metabolite variation (Corticosterone), total distance traveled (Distance), total roaming entropy (Entropy), total occurrences of defensive behaviors (Defensive), total occurrences of maintenance behaviors (Maintenance; drinking, eating, grooming), total occurrences of aggressive behaviors (Aggressive), total preference for the open area (Pref.open area), total occurrences of affiliative behaviors (Affiliative), area under the curve in the DDT (AUC.DDT), hub centrality in aggression network (HUB.agg), flexibility score in reversed-RGT (Flexibility), preference in last 20 min of rat gambling task (RGT), latency to collect pellet in RGT (Latency RGT), Blanchard dominance score (Blanchard). Panels A–B: +/+ n = 48, −/− n = 30.

## Discussion

In this multidimensional study, we used classical and ethological approaches of testing to evaluate the effects of brain serotonin deficiency on the expression of cognitive, social, and affective functions in different contexts and in the same animals. With unsupervised statistics, we identified which functions were primarily affected by the absence of brain serotonin. Surprisingly, no function evaluated in the classical testing appeared altered by its absence. However, in the day-to-day context of the home-cage, the absence of brain serotonin most strikingly affected the animals’ sexual, maintenance (eating, drinking, grooming), and defensive behaviors, levels of home-cage RE, weight, and corticosterone. These discriminative markers of serotonin function, consistent with the constellation of other behavioral impairments observed in *Tph2*^*−/−*^ rats, are reminiscent of common symptoms found in human impulse control disorders (ICD; e.g. disruptive, impulse control, and conduct disorders, compulsive sexual behavior disorder, and behavioral addictions) and stress and anxiety disorders (e.g. obsessive-compulsive, post-traumatic stress, and generalized anxiety disorders), which also share a high comorbidity level with ICDs [Table S10; (68–73)].

Under the complex and experimenter-free conditions of their home-cage, *Tph2*^*−/−*^ rats showed increased corticosterone levels, exacerbated repetitive aggression, and exploratory (sniffing) and sexual behaviors while neglecting affiliative (huddling), self-caring (grooming), and self-sustaining (feeding, poor maintenance of body weight) essential behaviors. While the dynamics of interactions eventually normalized for aggressive, exploratory, and affiliative behaviors, it did not for sexual behaviors. In clinical settings, cortisol disturbances, uncontrolled repetitive violent or sexual outbursts with poor consequences for others (harm) and self (neglect of health and personal care) are characteristic of disruptive, impulse control, and conduct disorders (74–78) and compulsive sexual behavior disorder (79). At the group level, *Tph2*^*−/−*^ hierarchical ranks appeared less stable and did not reflect in the structure of aggression networks (i.e. hub centrality) as was the case in the control groups. *Tph2*^*−/−*^ groups were disorganized overall; individuals presented a reactive coping style with persistent sexual activity and outbursts of aggression, appearing devoid of long-term goals (e.g. reproduction, secure food resource) and of specificity (e.g. occurred between random conspecifics). Additionally, *Tph2*^*−/−*^ rats expressed a hypervigilant defensive profile with higher day/night activity and smaller territories, ignoring food sources but favoring hiding and escaping options. Concerning the physiological changes, possible explanations could be that the downstream glucocorticoid receptor pathway’s disruption by serotonin depletion may have maintained elevated corticosterone levels in *Tph2*^*−/−*^ rats (80,81), and weight loss may have resulted from social stress-inducing feeding pattern modifications (82,83). Finally, the rich phenotype of the *Tph2*^*−/−*^ rats within the VBS confirmed the potential of this line to model transdiagnostic features of human disorders and revealed behavioral dysfunctions at the group level and the essential role of serotonin in modulating social and non-social daily life behaviors.

However, outside the home cage, the same animals had normal scores under the controlled conditions of cognitive testing. *Tph2*^*−/−*^ rats solved complex and risky decision-making tasks. They showed normal cognitive flexibility, typical sensitivity to reward, satisfactory motor control, good social recognition and odor discrimination abilities, and normal levels of anxiety and risk-taking. Only in the DDT, they appeared more sensitive to the discounting effect of the delay on their preference for the larger reward. Such preserved cognitive performance in the absence of brain serotonin was highly unexpected, as it contrasted with the dominant literature indicating an essential role of serotonin in modulating these higher-order functions using the same classical tests [(29,41,84–93), although see (94–102)]. However, before these results might indicate a more limited role for serotonin in modulating executive functions (decision-making, impulsivity, flexibility, social recognition), it is necessary to consider other potential explanations.

The lack of cognitive impairments could be due to the specific animal model we used. Knockout models specifically target one gene (103). Compared to pharmacological models, they prevent potential off-target effects associated with compound specificity, dosage, and application route. In a previous study, we confirmed normal cognitive and social abilities in *Tph2*^+/+^ rats (52), excluding the risk of a flooring effect in *Tph2*^*−/−*^ rats. However, a limitation of constitutive knockout models is their propensity to develop unexpected compensatory mechanisms, which might neutralize the genetic perturbation and result in a lack of phenotype (104). Following this hypothesis, *Tph2*^*−/−*^ rats have been found to show an increase in brain-derived neurotrophic factor (BDNF) levels in the hippocampus and prefrontal cortex (105– 107) and serotonergic hyperinnervation (107,108). Despite a complete lack of brain serotonin (96%; 108), *Tph2*^*−/−*^ mice present functional serotonergic neurons (110,111). Considering the physiological co-transmissions of glutamate, dopamine, or GABA neurotransmitters by serotonergic neurons, the activity of serotonergic circuitry could have occurred in the absence of serotonin (112–117). The hypothesis of such a compensatory scheme, counteracting the absence of brain serotonin in classical stand-alone cognitive tests, would suggest the existence of powerful biological targets for cognitive remediation, which remain to be studied.

Although it is unclear which compensatory mechanisms could have counterbalanced the absence of serotonin in classical tests, these mechanisms showed their limits under the less controlled, experimenter-free conditions of the social home-cage. In this more cognitively challenging and dynamic environment, *Tph2*^*−/−*^ rats presented altered daily life, social, and group behaviors compared to control rats. In classical tests, the cognitive demand is minimized to evaluating a few given functions, unlike natural environments where complex cognition is encouraged (56). Behavioral adaptation in social environments is known to be facilitated by serotonin through its influence on neural plasticity (30,118,119). Despite normal performances in classical cognitive tests, in the VBS, the highly dysfunctional social profile of *Tph2*^*−/−*^ rats indicates poor impulse control (e.g. sustained aggression), limited ability to adjust choices over time (e.g. sexual activity), and lack of goal-directed behavior (e.g. reduced eating and struggling at the feeder). Consistent with the context-specific role of central serotonin in modulating cognition (118,119), serotonin proved essential in supporting daily cognitive life in complex and social contexts.

Finally, an intriguing result concerns their social exploratory dynamic. Sniffing one another is a critical behavior in acquiring information (120), communicating dominance status (121), and pacifying interactions (122). *Tph2*^*−/−*^ rats showed slower reduction of sniffing network density in the VBS and a higher interest in the social partner in the social recognition test. They might be slower at integrating and transmitting social cues and thus at adjusting their behavior. The lack of structure of the aggression network may indicate disrupted transmission of hierarchical information in *Tph2*^*−/−*^ groups. Thus, communication deficits may have played a significant role in maintaining aggression, hierarchical disorganization, social stress, and the uncertainty level of the VBS, potentiating the serotonin depletion effects. A deeper investigation of the communication strategy of *Tph2*^*−/−*^ rats would help understand which functions affected by serotonin depletion are responsible for these deficits.

In this study, using adult *Tph2*^*−/−*^ rats, we showed that central serotonin was not essential for expressing cognitive abilities when tested in classical tests. However, central serotonin was a key modulator of essential naturalistic home-cage behaviors when living in undisturbed social groups. Context complexity must be integrated into experimental designs to investigate the role of the serotonergic system in the subtle modulation of different aspects of social and non-social behaviors. Only when facing the dynamic complexity and uncertainty of naturalistic conditions of choices were *Tph2*^*−/−*^ rats unable to adjust their behavior and were revealed as a promising model for studying transdiagnostic markers of ICDs and anxiety. The decision-making, flexibility, and impulsivity of the *Tph2*^*−/−*^ rats should be further studied under complex naturalistic conditions (123–125). In the complex social contexts, the unsupervised analysis of multidimensional results and analysis of network dynamics and hierarchy are essential additions to classical methods. They are necessary to expose the complexity of animals’ phenotypes and demonstrate the translational value of results.

## Supporting information

ARRIVE Checklist

## Acknowledgments

This work was funded by a DFG grant (RI 2474/2-1 to Marion Rivalan and AL 1197/5-1 to Natalia Alenina) and supported by project 73022475 of the St. Petersburg State University to Natalia Alenina. Support was also received through DFG funding to the Center of Excellence NeuroCure DFG EXC 257. We want to thank Nina Soto, Vladislav Nachev, Patrik Bey, Miléna Brunet, Tania Fernández del Valle Alquicira, Franca Eltner, Arnau Ramos-Prats, Diane Delgado, Melissa Long, Alexej Schatz, Martin Dehnhard, and his team for their help with the experiments, data extraction, and analysis. We thank Annegret Dahlke, Monique Bergemann, Laura Rosenzweig, Susanne da Costa Goncalves, Bettina Müller, Reimunde Hellwig-Träger, and the FEM team for their technical assistance with the animals. We thank our colleagues in the Winter lab who made insightful comments on a previous version of the manuscript.

## Declaration of interest

YW owns PhenoSys equity. All other authors declare no conflict of interest.

## Notes

https://doi.org/10.5281/zenodo.4912528

