## Supplemental information for "Constitutive depletion of brain serotonin differentially affects rats’ social and cognitive abilities"

### 1. SUPPLEMENTAL METHODS

#### A. Animals and housing conditions

The *Tph2*$^{+/+}$ group consisted of 10 Dark Agouti and 38 TPH2-ZFN rats (1). All animals were born at the Max Delbrück Center for Molecular Medicine, Berlin. Animals arrived at our animal facility between five and nine weeks of age, they were housed in standard rat cages (Eurostandard Type IV, 38 cm x 59 cm). We used 8 *Tph2*$^{+/+}$ and 5 *Tph2*$^{-/-}$ cohorts, 6 animals each. Groups of 12 animals (6 *Tph2*$^{+/+}$ and 6 *Tph2*$^{-/-}$) were tested either in the morning or in the afternoon (i.e. 24 animals per day) depending on the light cycle of the housing room (lights on at 20:00 in room 1 or 01:00 in room 2) in order to maximize the use of our four operant cages and minimize potential circadian effect (rats were all tested in RGT within 3h and 1h after start of dark phase). Animals were fed with standard maintenance food (V1534-000, Ssniff, Germany). Numbers of animals for each test are reported in the Table S1. The number of animals was decided following *a priori* power analysis (n = 51, Gpower 3.1.2). It was reduced because of the difficulty of breeding of TPH2-ZFN rats and the low survival rate of *Tph2*$^{-/-}$ rats.

#### B. Handling

After staying a week undisturbed in the animal facility, animals were handled daily by the experimenters. Since *Tph2*$^{-/-}$ animals were very reactive to manual handling all animals were handled using a 6 cm diameter grey polypropylene tube that was added in the cage as enrichment and used by the animals as shelter preventing fights and mounting behavior.

#### C. Marking

Two weeks before the beginning of the training phase, rats were marked individually, subcutaneously in the ventral left lower quadrant with a radio-frequency identification (RFID) chip (glass transponder 3x13 mm, Euro I.D.) under short isoflurane anesthesia.

#### D. Testing

The order of the tests and inter-test pauses were chosen to minimize any interference of one test on another. Training and testing started 1 h after the beginning of the dark phase. Animals were habituated to the experimental room conditions for 30 min before the start of the test. The order of testing of the animals was mixed and balanced in order to minimize potential confounders. A randomly generated sequence was not used for that. Blinding of the experimenter to the genotype of the animals was not possible during the conduct of experiment due to important behavioral differences at baseline. Automatic outcome assessment was used for data collection for all tests except dark light box test, social recognition task, odor discrimination test and video scoring of the visible burrow system test.

##### E. Operant system

A clear partition with a central opening in the middle of the operant cage ensured an equal distance to all nose-poke holes from this central opening for an approaching rat.

##### F. Rat Gambling Task (RGT)

The operant cages were equipped with four nose-poke holes on the operant wall.

The training 1 started with the four nose-pokes lit and active, a single nose-poke generated the delivery of one pellet. The selected hole remained lit until the collection of the pellet into the magazine while all the other holes were inactive. A visit to the magazine induced the reactivation and illumination of all the nose-poke holes. The training 1 continued daily until rats obtained 100 pellets in a session (30 minutes cut-off), then they could start the training 2. In training 2, two consecutive nose-pokes at the same hole were required to obtain one pellet and the same criterion had to be reached (100 pellets in 30 minutes cut-off). In training 3, two pellets were delivered after a choice (two consecutive nose-pokes) during a short session (maximum 30 pellets and 15 minutes cut-off). A forced training (2) was applied to counter any side preference developed during the training procedure: if the choices for the two holes of one side were superior to 60% during the last session of training 2. During the first part of the forced-training, the two nose-poke holes on the non-preferred side were active and lit, two consecutive nose-pokes into the active holes induced the delivery of one pellet. The holes on the preferred side were inactive and not lit. After the collection of 15 pellets, the second part of the forced training started with the four holes active and lit. Two consecutive nose-pokes into holes of the preferred side induced the delivery of one pellet with a probability of 20% whereas choosing the non-preferred side induced the delivery of one pellet with a probability of 80%. The cut-off was 50 pellets or 30 minutes. The training procedure lasted six to ten days and the test was performed the next day.

During the test, each of the four holes was associated to an amount of reward and a possible penalty (time-out) which was unknown to the rat. Two holes on one side were rewarded by two pellets and associated to unpredictable long time-outs (222s and 444s with the probability of occurrence ½ and ¼ respectively), in the long term those options were disadvantageous. The two holes on the other side were rewarded by one pellet and associated to unpredictable short time-outs (6s and 12s with the probability of occurrence ½ and ¼ respectively), in the long term those options were advantageous. After a choice (two consecutive nose-pokes), the reward was delivered and the selected hole remained lit until a visit to the magazine or the duration of the time-out. During this time all the nose-poke holes were inactive. The test lasted one hour (or cut-off 250 pellets). The theoretical maximum gain of the advantageous options was five times higher than the disadvantageous options at the end of the test (60 min). The percentage of advantageous choices for the last 20 min of RGT was used to identify good decision-makers (GDMs) >70% of advantageous choices, poor decision-makers (PDMs) <30% of advantageous choices and intermediate animals. The percentage of advantageous choices per ten minutes indicated the progression of the preference over time. An index of the motivation for the reward was measured as the mean latency to visit the feeder after a choice.

##### G. Reversed-RGT

The animals were tested in the reversed-RGT (2,3) 48 hours after the RGT. The same advantageous and disadvantageous options as in the RGT were used but they were switched from one side to the other. The test lasted one hour (or cut-off 250 pellets).

A flexibility score was calculated as the preference for the location of the non-preferred option during the RGT. Flexible rats had >60% of such choices during the last 20 minutes, undecided rats had between 40% and 60% of choices, and inflexible rats had <40%.

**H. Delay discounting task (DDT)**

The operant cages were equipped with two nose-poke holes the furthest from each other on the operant wall (2,3). One nose-poke hole (NP1) was associated with a small immediate reward (1 pellet) and a second nose-poke hole (NP5; 25 cm between the two holes) with a large (5 pellets) reward. During the training, the large reward was obtained immediately (delay 0sec) after the choice (two consecutive nose-pokes). After the pellet delivery, the magazine and house lights were turned on for a 60s time-out. The session lasted 30 minutes (or cut-off 100 pellets). A percentage of choice of the large reward ≥70% on two following sessions with ≤15% variation (stability criterion) was required to start the test. Minimum three training sessions were done. During the test, choosing NP5 induced the delivery of the large reward after a designated delay, NP5 stayed lit during the duration of the delay. After the pellet delivery of the large reward the magazine and the house lights were turned on for a time-out of 60s minus the duration of the delay. The delay was fixed for a day and increased by 10s from 0s to 40s according to a stability criterion ≤10% variation of choice of the large reward during two consecutive sessions. The test sessions lasted 60 minutes (or cut-off 100 pellets).

The preference for the large delayed reward was calculated as the mean percentage of NP5 choices during two stable sessions. To calculate the area under the curve (AUC) which represents the sensitivity to delay, for each individual the preference for the large delayed reward for each delays was normalized to the preference for the large delayed reward during the training and plotted against delay as a proportion of maximum delay (4); the area under this normalized curved was then calculated.

##### I. Probability discounting task (PDT)

The operant cages were equipped with two nose-poke holes the furthest from each other on the operant wall. It is an adaptation from the test described in Alonso et al., 2019 (2) with the addition of a stability criterion. During the training, the large reward was always delivered after choosing NP5 (probability P=1), which allowed the rats to develop a preference for NP5. Two consecutive nose-pokes induced the delivery of the reward after 4s, during this time the selected hole stayed lit. Then the magazine light turned on for a 15s time-out. The session lasted 25 minutes (or cut-off 100 pellets). A percentage of choice of the large reward ≥70% on two following sessions with ≤15% variation (stability criterion) was required to start the test. At least three training sessions were done. During the test, the probability we used P = 0.66, 0.33, 0.20, 0.14 and 0.09. The probability was fixed for a day and increased according to a stability criterion ≤10% variation of choice of the large reward during two consecutive sessions. The session lasted 25 minutes (or cut-off 100 pellets).

The percentage of preference for the large and uncertain reward was calculated for each probability as the percentage of NP5 choices during the two stable sessions. To calculate the AUC which represents the sensitivity to probabilistic uncertainty and risk taking, for each individual the preference for the large reward for each probability was normalized to the preference for the large reward during training and plotted against probabilities expressed as odds (5) with odds = (1/P) -1; the area under this normalized curved was then calculated.

##### J. Fixed-interval and extinction schedule of reinforcement test (FIEXT)

The operant cages were equipped with a central single nose-poke hole or a single lever. The fixed-interval, FI, consists of two phases: a fixed time interval during which choices are not rewarded, followed by a phase where a choice can be rewarded (3). The extinction, EXT, is a longer, fixed time interval during which no choices are rewarded. Both FI and EXT are conditions that cause frustration in the animal. A session consisted of the repetition of seven FI and one EXT of 5 min. The maximum number of pellets was 14 during a single session. FI lasted 30 s for the first four sessions, 1 min for the next four sessions, 2 min for the next three sessions and 1 min for the final four sessions. The final four sessions with a 1 min FI were the actual test. During the FI, the house light was on and the central nose-poke hole was inactive. At the end of the FI, the house light turned off and the central nose-poke was lit and became active; two consecutive nose-pokes induced the delivery of one pellet, the central nose-poke light was turned off and the tray light was lit. A visit to the tray induced the start of the next FI. After seven consecutive FI, the EXT period started, with all lights off and no consequences associated with nose poking.

When the operant cages were equipped with a lever, the scheme was similar. During the FI, the house light was on and any press on the lever had no consequence. At the end of the FI, a cue light above the lever turned on and the first press was rewarded by a pellet. The cue light above the lever stayed on until pellet collection. A visit to the tray induced the start of the next FI. After seven repetitions of the FI and pellet collection the EXT started. During EXT the house light was off and any press on the lever had no consequence.

As described earlier (6), the data from the first FI of the session and the first FI after the first EXT were excluded. The total number of nose pokes and mean number of nose pokes were determined for each FI and EXT period. We summed nose pokes for 10 s intervals during FI to visualize the anticipatory activity of the rats. Likewise, we summed nose pokes for 1 min intervals during EXT to visualize the perseverative activity.

##### K. Social recognition task (SRt)

This test was adapted from Shahar-Gold et al., 2013 (7) and is described in Alonso et al., 2019 (2). The test took place in a square open field (OF, 50 x 50 cm), a small cage was placed in one corner of the OF. To improve the setup, a foam PVC partition was placed around this intruder's cage to avoid the test rat hiding behind the cage. The unfamiliar conspecifics were older Wistar Han rats, accustomed to the procedure. A video camera on top of the OF recorded the experiment. Each rat was tested on two consecutive days. On the first day, the subject was placed in the OF containing the empty cage in a corner for a habituation of 15 minutes. Then, the unfamiliar conspecific was place in the small cage and the subject was allowed to freely explore the open field for five minutes (E1). After that the small cage with the conspecific was removed from the open field, and the subject remained alone in the open field for a break of 10 minutes. The encounter procedure was repeated two more times with the same conspecific (E2, E3). On the second day, the first 15 minutes habituation phase was followed by a 4th encounter (E4) of five minutes encounter with the same conspecific as in day 1. After this encounter, a break of 30 minutes took place, in which the subject remained alone in the open field. Then, the last encounter took place, but a new unfamiliar conspecific was placed in the same small cage for five minutes (Enew).

The time spent in close interaction with the intruder was measured for each encounter and for the first five minutes of Habituation (Hab) when the subject smelled at the grid of the empty cage. The social preference was calculated as the ratio of the interaction time in E1 and Hab. The short-term social recognition was calculated as the ratio of the interaction time in E1 and E3. The long-term social recognition was calculated as the ratio of the interaction time in E4 and Enew.

#### L. Odor discrimination test

The test took place in a square OF (50 x 50 cm). Two plastic petri dishes filled with either spoiled or fresh bedding were placed in two opposite corners of the OF. A video camera on top of the OF recorded the experiment. The test rat explored the OF for 5 min. The time spent in close interaction with each dish was measured and the preference for the spoiled bedding (social odor) was calculated.

**M. Dark light box test**

A box with two compartments of 45 cm x 22.5 cm x 35 cm, one bright compartment made of transparent plastic and one dark compartment made of black opaque plastic and with a lid of the same material. A gate (9 cm x 10 cm) enabled the rats to pass from one compartment to the other. Room light was on and extra lamps were positioned above the box providing a high light intensity in the bright compartment > 500 lux. Inside the dark box there was no appreciable illumination (i.e., 2 lux). The rat was brought into the bright compartment (with the home-cage tube) and allowed to explore the apparatus for 10 min. After the test, the apparatus was cleaned with 5% ethanol before the next rat was assessed. We recorded each tests with a video camera placed above the bright compartment. We measured the number and duration of visits to each compartment, number of risk assessments which included head poking through the door and body stretches, the latency to leave the bright compartment the first time and the duration of the first visit to the dark compartment. Risk taking index (8) was calculated as the sum of the duration of the first visit to the dark compartment, the number of risk assessment into the light compartment and the time spent in the dark compartment, for clarity this number was subtracted to the maximum score in order to get ascending values.

**N. Automated visible burrow system (VBS)**

The automated VBS, as previously described (2) consisted of an open area connected through two transparent tunnels to a burrow system made of infrared-transparent black plastic that remained in the dark throughout the test. Food and water were available at all time in the open area. The burrow system consisted of a large chamber, a small chamber and a tunnel system. A grid of 32 RFID detectors was placed underneath the VBS in order to automatically determine individual animal positions using the program PhenoSoft (PhenoSys, Berlin). An infrared camera (IP-Camera NC-230WF HD 720p, TriVision Tech, USA) mounted above the VBS recorded a 30 s video every 10 min (CamUniversal, CrazyPixels, Germany). The software PhenoSoft ColonyCage (PhenoSys, Berlin) was used to identify individuals in the videos. Six rats of the same genotype were housed in the VBS for seven days in a humidity- and temperature-controlled room (temperature 23-24°C, humidity 45-50%) containing two VBS systems. The animals were visually checked every day. After the first group (6 *Tph2*$^{+/+}$ and 6 *Tph2*$^{-/-}$), the duration of the VBS housing was reduced from seven to four days (9) for the *Tph2*$^{-/-}$ animals due to noticeable weight loss.

The videos of the first four hours of the dark and light phases were scored using a scan sampling method (63). All aggressive behaviors except "struggling at feeder" were grouped under "general aggression" and sexual behaviors grouped under "sexual". We present the most expressed behaviors (median > 5): huddling, sniffing, eating, grooming, general aggression, struggling at feeder and sexual. All scored behaviors (Table 1 of the main text) are shown in the figure S5. The body weight of the animals was measured before and after VBS housing (4 or 7 days); the difference of weight was calculated. Although wounds were rarely observed during this study, they were documented at the end of VBS housing. The activity (distance traveled) and the place preference were extracted using the software PhenoSoft analytics (PhenoSys, Berlin) for the first four days of VBS housing. The time spent in the open area of the VBS was measured using the data collected from the grid of detectors.

##### O. Glicko rating system

Social ranking of the rats was determined using the Glicko rating system based on aggressive behaviors during the dark phase (14). The Glicko rating (10,11) was calculated within each VBS groups using the R package 'PlayerRating' (12). The direction of the interaction defined the winning animal (initiator) and losing animal (receiver). We considered all types of aggressive and sexual behaviors for the first four days because sexual behaviours elicited defensive behaviors and vocalizations in the receiver. The Glicko rating is a dynamic measure that updates for each individual after each interaction. We used the R package 'online CPD' (13) to detect the change points of the Glicko rating over time for each individuals and determine the stability of the rating. Because the total number of agonistic interactions varied between VBS groups, we calculated a normalized number of change points dividing by the group total number of interactions. For each group the divergence or maximum rating contrast was the difference between the highest and the lowest individual final ratings. Dominant animals' ratings were higher than 1/3 of the maximum rating contrast of the group.

##### P. Blanchard dominance score

The Blanchard dominance score (14) is a dominance score established in the original VBS. It originally combines three classical parameters: the number and location of wounds, the time spent in the open area and the weight loss. A wound is a visible alteration of the skin of an animal such as scratches and scabs. A wounded animal was monitored closely until complete skin healing. In our study wounds rarely occurred. Over the 78 rats tested, only nine rats presented one to 6 wounds in total (over 4 to 7 days in VBS). Because of its sporadic occurrence, the number of wounds could not be considered in the calculation of the Blanchard dominance score. For each individual within a group, time spent in open area and weight loss for the entire stay in the VBS (7 or 4 days) were ranked from 1 to 6, the average of both ranks was the Blanchard dominance score.

##### Q. Roaming entropy

The roaming entropy (RE) within the VBS, is the probability for an individual to be at a certain place at a given time. RE calculation was based on the method described in Freund et al., (63). In the automated VBS, it indicates the spatial dispersion of the rats. Continuous location recordings from the RFID grid were cleaned and filtered; we selected the data from the dark phase of the first four days. We sliced the data into 1 s detections for each rat in order to weigh longer detections. We calculated the observed frequencies or probabilities, $p_{i,j,d}$ of detection of each animal $i$ at each reader $j$ on a day $d$. These frequencies were then used to compute the RE for each day, following the equation of Shannon: $RE_{i,d} = - \Sigma (p_{i,j,d} \log p_{i,j,d}) / \log(k)$ where k is the number of detectors in the automated VBS. The total roaming entropy was calculated for the four days.

**R. Social Network Analysis (SNA)**

We developed the method to social network analysis to understand the qualitative aspects of the social interactions between the individuals. It allows uncovering individual and group dynamics such as information transmission or power distribution. Behavioral interactions between two individuals were organized into matrices for each category of behavior (huddling, sniffing, struggling at feeder, aggression, and sexual behavior). The matrices were weighted and directed, meaning that the number of occurrences of interactions was used and that all interactions weren't always reciprocal in a pair of rats. We used the R package igraph (15) to calculate the parameters and visualize the networks.

We measured three global network parameters: density, average path length and out-degree centralization to understand the structure of the networks (11). Density is the proportion of possible ties that can exist in the network. Average path length is the mean number of steps between any pair of individuals in the network. Out-degree centralization indicates the differences of initiated connections between the individuals. We measured five individual network parameters: in- and out-degree, betweenness centrality, closeness centrality, Bonacich's power centrality and Hub centrality, to understand the roles of individuals within networks (11). In- and out-degree is the number of interactions an individual receives and initiates respectively. Betweenness centrality indicates how much an individual connects two other individuals. Closeness centrality indicates how much an individual directly connects with other individuals. Bonacich's Power Centrality defines the influence of an individual based on the connections of its neighbors, powerful individuals are connected to many individuals that themselves are less connected to others (16). Hub centrality also depends on the connection of an individual's neighbors, powerful individuals (authorities) are connected to many individuals highly connected to others (hubs) (17).

##### S. Feces collection and corticosterone metabolites measurements

One day before and immediately after the VBS stay, both times at the same time of the day, the rats were housed in individual cages with food, water and clean bedding for 4 hours maximum. Every 30 minutes, feces produced were collected in microtubes and stored at -20°C until extraction. Then, the samples were defrozen, 0.1g of feces was added to 0.9 ml of 90% methanol, agitated for 30 minutes and then centrifuged at 3000 rpm for 15 minutes. A 0.5 ml aliquot of the supernatant was added to 0.5 ml of water, this extract was stored at -20°C. Corticosterone metabolite measurements were performed with enzyme immunoassay (EIA) following the method of (18) in the laboratory of Dr. Dehnhard at the Leibniz Institute of Zoo and Wildlife Research. Briefly, a double antibody technique was used in association with a peroxidase conjugate generating a signal quantitatively measurable by photometry. Concentration expressed in micrograms/grams of feces.

##### T. Statistical analysis

Before comparing the genotypes, we compared the performance of Dark Agouti (n=10) and Tph2-ZFN (n=38) animals in all the tests using the Wilcoxon rank sum test. The results from the Dark Agouti and Tph2-ZFN control groups were not different and grouped together to form the Tph2-control group ($Tph2^{+/+}$). During the data analysis the experimenter was not blind to the genotype of the animals. We do not expect our data to follow a normal distribution; hence we used non-parametric statistical tests. We used the Wilcoxon rank sum test to compare the two genotypes ($Tph2^{+/+}$ *vs.* $Tph2^{-/-}$) against each other, the Fisher's exact test to compare the number of GDMs and PDMs in *Tph2*$^{+/+}$ and *Tph2*$^{-/-}$ groups, the one sample t-test to compare the performance of the animals to a theoretical value in RGT and the Wilcoxon sign test [R package RVAideMemoire, (19)] to compare the performance of the animals to a theoretical value in DDT, PDT, SRt and odor discrimination test. Differences in performance between GDMs and PDMs were evaluated with the cohen's effect size [R package effsize, (20)]. Linear mixed-effect models (lmer models) can be used robustly on non-normal data (21). We used lmer models [R package lmerTest, (22)] to compare genotypes (or decision maker groups) over several time points and with individual and batch information as nested random effects. Post-hoc multiples comparisons were done on the linear models [R package multcomp, (23)], there the p-values were adjusted using the Holms method for multiple comparisons. Because of their ability to model over-dispersion, we used generalized linear models with Markov chains [MCMCglmm, R package MCMCglmm, (24)] to compare the distance traveled of genotypes over light cycles and hours with individual and batch information as random effects. The fitting of the MCMCglmm models was assessed with the plots of the fixed effects and random effects. The lower deviance information criterion (DIC) was used to choose the best MCMCglmm model. We used Spearman's correlation [R package Hmisc, (25)] to assess the link between hierarchy variables (Glicko and Blanchard scores), individual SNA centrality, roaming entropy and corticosterone level after VBS stay. The Random Forest [RF, R package randomForest, (26)] predicts the genotype of each individual based on their scores in each test and returns the importance of each variable for the classification. We used a Leave-One-Out cross-validation and ran the RF for 100 runs. A k-means clustering [R package stats, (27)] grouped the variables by importance; the number of clusters (n = 4) was chosen to maximize homogeneity within a cluster and minimize homogeneity between clusters (Fig. 5B). The Principal Component Analysis [PCA, R package stats, (27)] summarizes the data set in new dimensions representing which is the most variable between individuals. RF and PCA were run on the same datasets. As both methods cannot handle missing data; they were run on a selection of variables including all animals of the study, and additionally on two other sets with more variables but excluding some groups of animals (see Table S5, S6, S7 and S8).

### 2. SUPPLEMENTAL FIGURES

#### A. Figure S1: Progression of choices per type of decision maker

Decision makers of both genotypes showed similar progression of choices (Fig. S1).

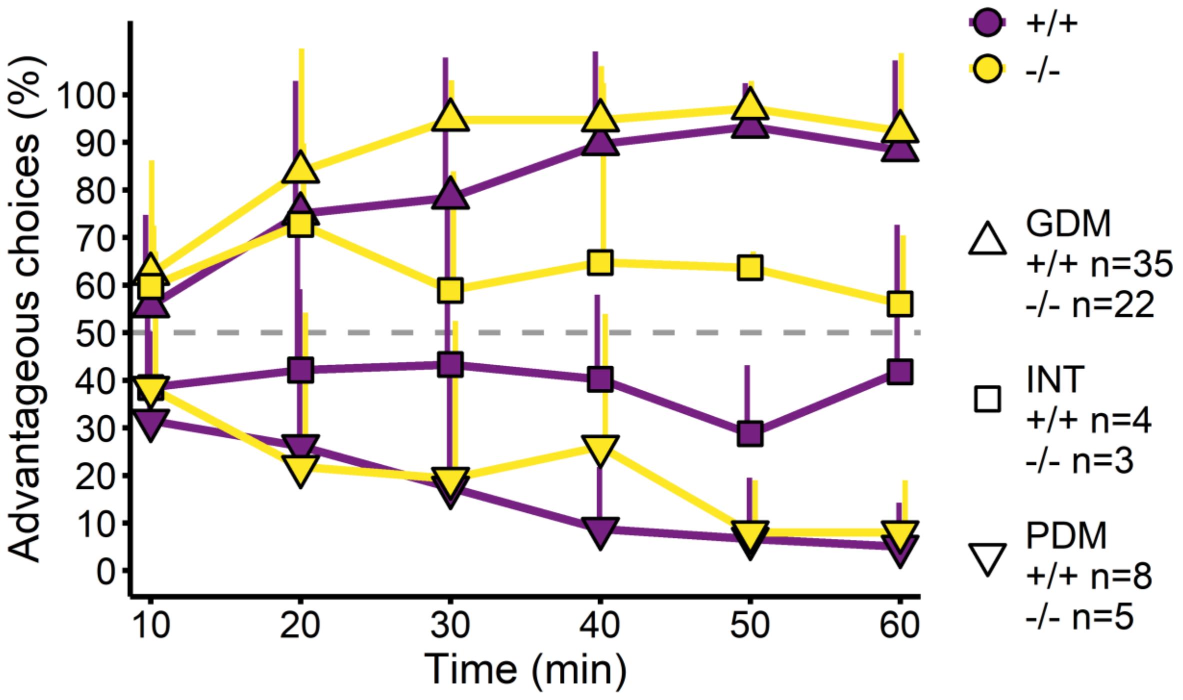

**Figure S1. Progression of choices per type of decision maker.** Good decision makers (GDM. upward triangle). intermediates (INT. square). and poor decision makers (PDM. downward triangle). Lines indicate mean + SD. *Tph2*$^{+/+}$ in purple and *Tph2*$^{-/-}$ in yellow.

#### B. Figure S2: Responses in FIEXT task with lever or nose-poke manipulandum

Anticipatory and perseverative behaviors were influenced by the manipulandum available in the operant cage. When a lever was used to express the behavior, *Tph2*$^{+/+}$ and *Tph2*$^{-/-}$ were not different across time and in total (Fig. S2A-left, S2B-left, S2C-left and S2D-left). However, when a nose-poke hole was used to express the behavior, *Tph2*$^{-/-}$ expressed less anticipatory behavior than *Tph2*$^{+/+}$ (Fig. S2A- right, Wilcoxon rank sum test, W = 269, p-value = 0.012) and less perseverative responses than *Tph2*$^{-/-}$ (Fig. S2B- right, Wilcoxon rank sum test, W = 275, p-value = 0.008). The comparison of the manipulandum inside each genotype showed that *Tph2*$^{-/-}$ expressed less anticipatory responses with the nose-poke hole than with the lever (Fig. S2C, Wilcoxon rank sum test, W = 120, p-value = 0.004) and that *Tph2*$^{+/+}$ expressed more perseverative responses with the lever than with nose-poke hole (Fig. S2D, Wilcoxon rank sum test, W = 105.5, p-value = 0.039).

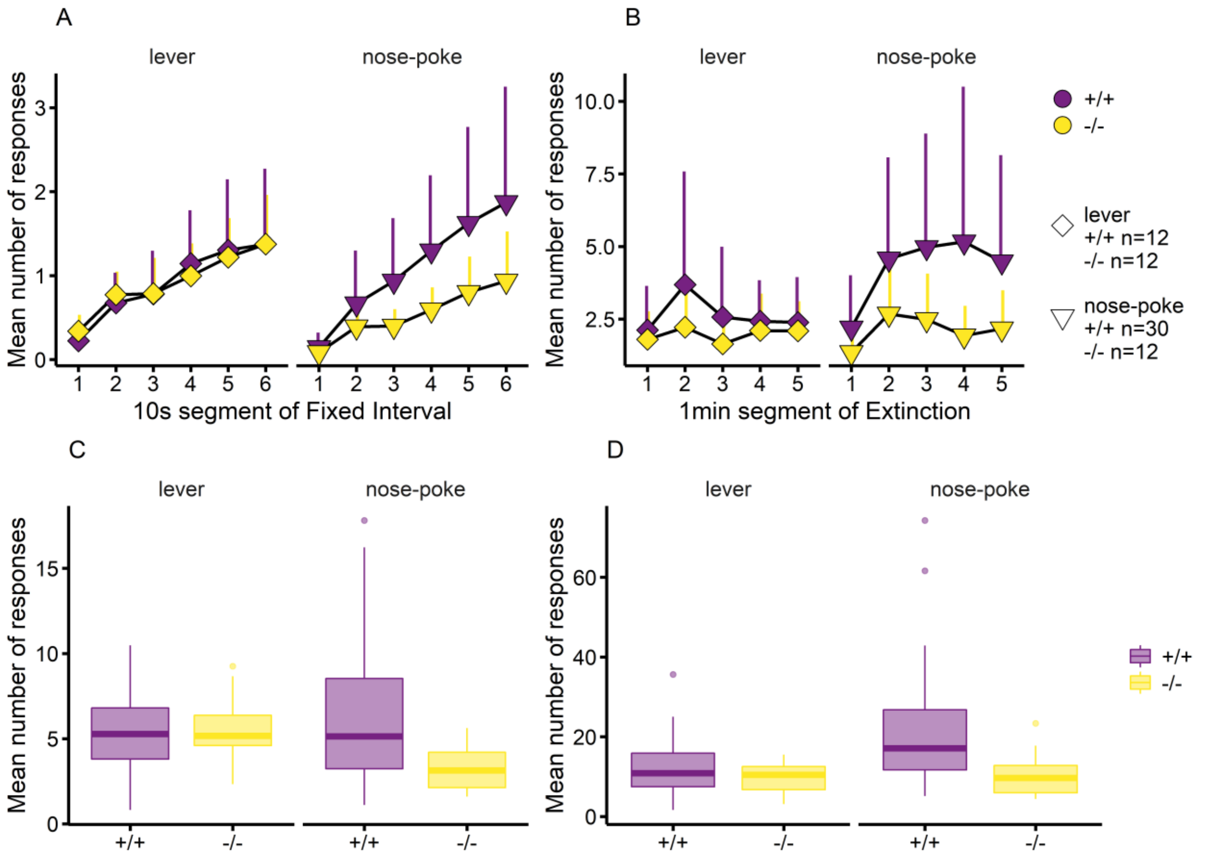

**Figure S2. Responses in FIEXT task with lever or nose-poke manipulandum A-**Mean number of responses during FI per 10s, lines indicate mean + SD. **B-**Mean number of responses during EXT per min, lines indicate mean + SD. **C-**Mean number of responses in FI. **D-**Mean number of responses in EXT. *Tph2*$^{+/+}$ in purple and *Tph2*$^{-/-}$ in yellow

##### C. Figure S3: Social recognition task ratios

*Tph2*$^{+/+}$ and *Tph2*$^{-/-}$ expressed similar social preference for the unfamiliar intruder during the first encounter (Fig. S3A, Wilcoxon rank sum test with continuity correction, W = 417.5, p-value = 0.6361). They expressed similar short-term social recognition over encounters (Fig. S3B, Wilcoxon rank sum test, W = 423, p-value = 0.6973). Long-term social recognition was not witnessed on the next day in both genotypes (Fig. S3C, ratio equal to 1 for both groups, Wilcoxon rank sum test, W = 439, p-value = 0.958). Absence of long term social recognition in control groups prevents possible interpretation of this phase of the test.

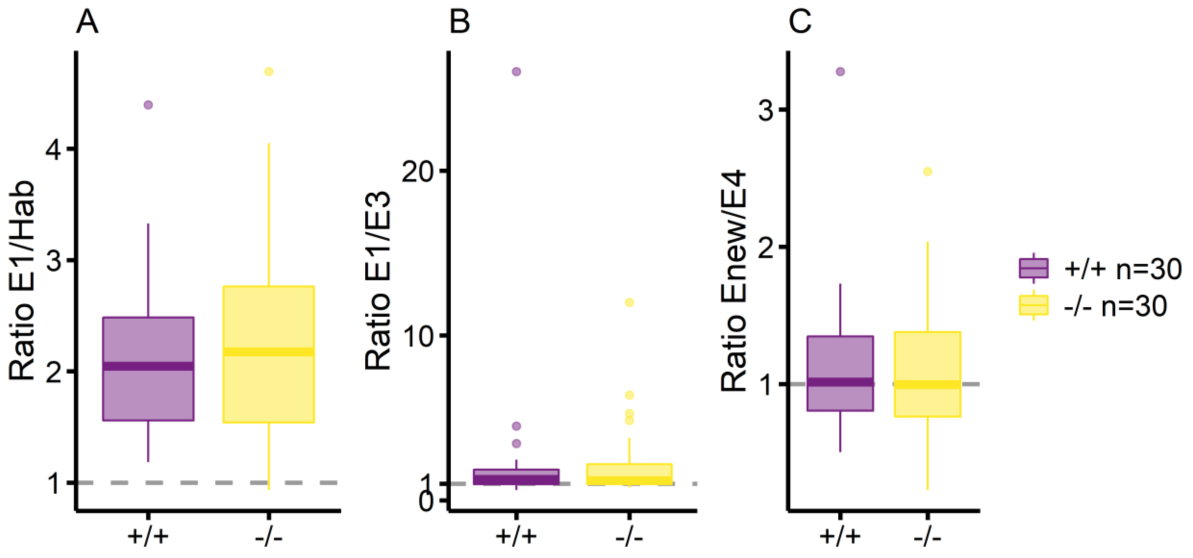

**Figure S3. Social recognition task ratios. A-**Social preference calculated as the ratio of the interaction time in E1 and habituation (hab) with and without the intruder. **B-**Short term social recognition memory calculated as the ratio of the interaction time in E1 and in E3 with the same intruder. **C-**Long term social recognition memory calculated as the ratio of the interaction time in Enew and in E4 with a new intruder *versus* the familiar intruder as in E1, E2, and E3. *Tph2*$^{+/+}$ in purple and *Tph2*$^{-/-}$ in yellow

##### D. Figure S4: Preference for the social odor in the odor discrimination test.

Both genotype showed a preference for the dish with the social odor was higher than chance (Fig. S4, Wilcoxon sign test, +/+: 95CI [57, 71], p-value < 0.001; -/-: 95CI [50, 76], p-value = 0.034) and at similar level (Fig. S4, Wilcoxon rank sum test with continuity correction, W = 330, p-value = 0.3921).

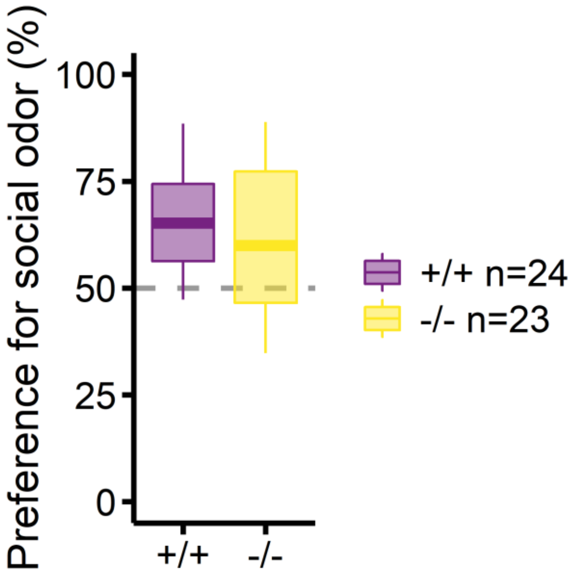

**Figure S4. Preference for the social odor in the odor discrimination test.** *Tph2*$^{+/+}$ in purple and *Tph2*$^{-/-}$ in yellow

#### E. Figure S5: All behaviors expressed in the VBS

Concerning the affiliative behaviors, *Tph2*$^{+/+}$ expressed more huddling (Wilcoxon rank sum test with continuity correction, W = 1240.5, p-value = 9.172e-08) and allogrooming (Wilcoxon rank sum test with continuity correction, W = 1086, p-value = 9.623e-05) than *Tph2*$^{-/-}$ rats. *Tph2*$^{-/-}$ rats expressed more sniffing directed to the nose than *Tph2*$^{+/+}$ rats (Wilcoxon rank sum test with continuity correction, W = 315.5, p-value = 3.177e-05). Concerning the maintenance behaviors, *Tph2*$^{+/+}$ expressed more eating (Wilcoxon rank sum test with continuity correction, W = 1267, p-value = 1.962e-08) and grooming (Wilcoxon rank sum test with continuity correction, W = 914.5, p-value = 0.04591) than *Tph2*$^{-/-}$ rats. Concerning the aggressive behaviors, *Tph2*$^{+/+}$ expressed more struggling at feeder (Wilcoxon rank sum test with continuity correction, W = 1227.5, p-value = 1.81e-07) than *Tph2*$^{-/-}$ rats. For all other aggressive behaviors, *Tph2*$^{-/-}$ animals had a higher number of occurrences than *Tph2*$^{+/+}$ animals (Wilcoxon rank sum test with continuity correction, struggling in tunnel: W = 233.5, p-value = 5.31e-07, mutual upright posture: W = 118.5, p-value = 5.426e-10, pinning: W = 268, p-value = 2.66e-06, fight: W = 452, p-value = 0.003342, attack: W = 518, p-value = 0.004951, following: W = 485, p-value = 0.001977, aggressive grooming: W = 361, p-value = 2.963e-05, except for struggling at water n.s.). Concerning the sexual behaviors, *Tph2*$^{-/-}$ rats performed more embracing (Wilcoxon rank sum test with continuity correction, W = 88.5, p-value = 1.198e-11) and mounting (Wilcoxon rank sum test with continuity correction, W = 113, p-value = 3.823e-13) than *Tph2*$^{+/+}$ rats.

Concerning the defensive behaviors, the occurrences were rare in both groups, nevertheless *Tph2*$^{-/-}$ rats performed more defensive behaviors in total (Wilcoxon rank sum test with continuity correction, W = 14166, p-value = 2.977e-06).

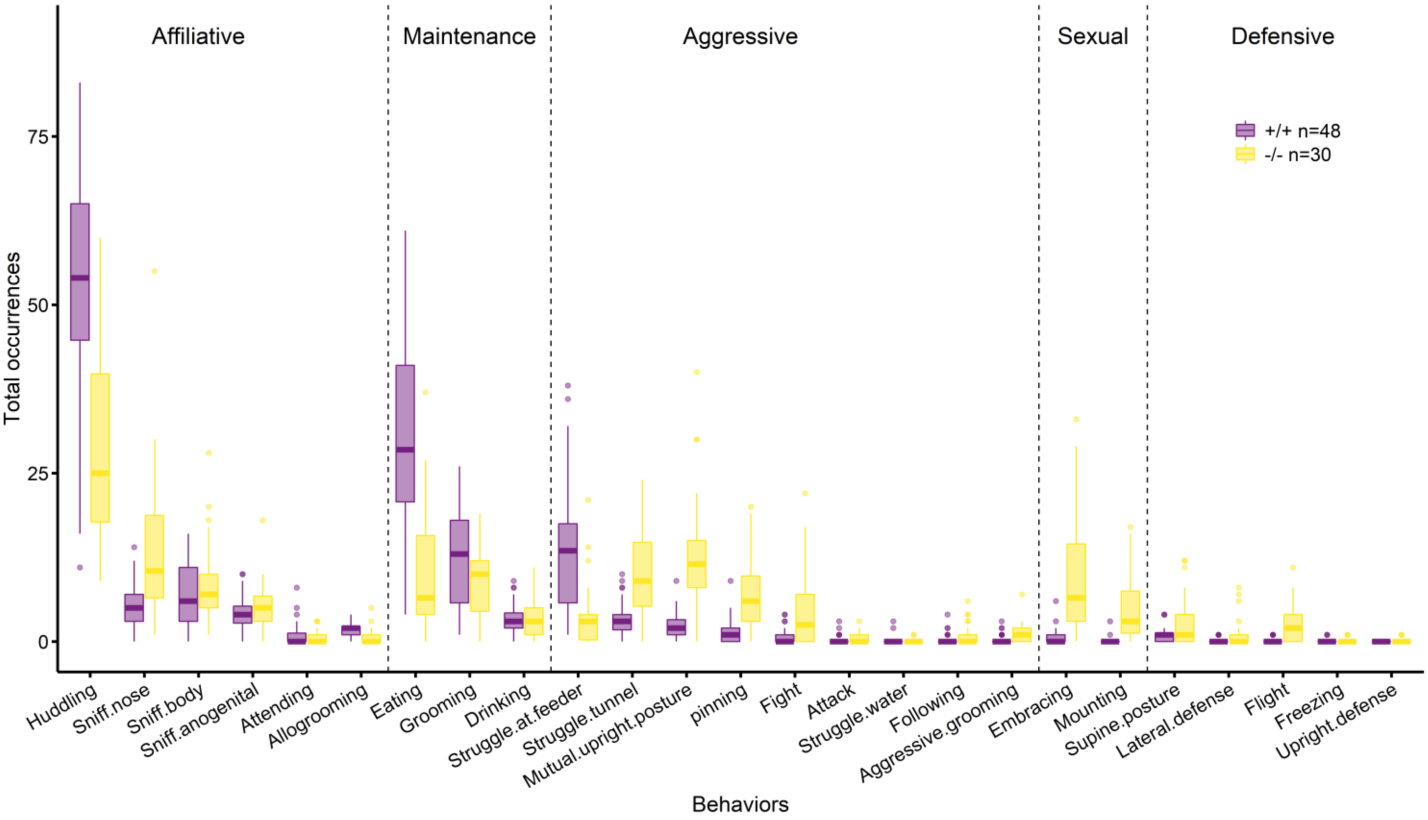

**Figure S5. All behaviors expressed in the VBS.** Vertical dash lines separate the categories of behaviors. *Tph2*$^{+/+}$ in purple and *Tph2*$^{-/-}$ in yellow.

##### F. Figure S6: Average path length and out-degree centralization of the social networks

Over time the average path length between two individuals reflected the density of the network. In huddling networks where the *Tph2*$^{-/-}$ groups became more closely connected, the average path length decreased [Fig. S6A, lmer *day*: $F(3,14) = 4$. p-value = 0.0208] and in "general aggression" networks where the *Tph2*$^{-/-}$ groups became less closely connected, the average path length increased [Fig. S6A, lmer *day*: $F(3,10) = 13$, p-value < 0.001]. The average path length of the sniffing networks was lower in *Tph2*$^{-/-}$ groups than control groups [Fig. S6A, lmer, *genotype*: $F(1,43) = 5$, p-value = 0.0319] with the higher density this indicated a higher cohesion of the sniffing networks in *Tph2*$^{-/-}$ groups. For struggling at feeder and sexual behaviors the average path length was not different neither between genotypes or days (Fig S6A, high variance). Finally, we looked at the out-degree centralization for which high values indicate differences in power between the individuals of the groups. Overall out-degree centralization values were low indicating that the distribution of power in sniffing, huddling, struggle at feeder, general aggression and sexual behaviors was relatively well distributed between individuals of the groups. Still, *Tph2*$^{-/-}$ groups had a lower centralization in struggling at feeder networks and higher centralization in sexual behavior networks than *Tph2*$^{+/+}$ groups [Fig. S6B, lmer, SAF: *genotype*: $F(1,44) = 8$, p-value = 0.0072; Sexual: *genotype*: $F(1,41) = 29$, p-value < 0.001].

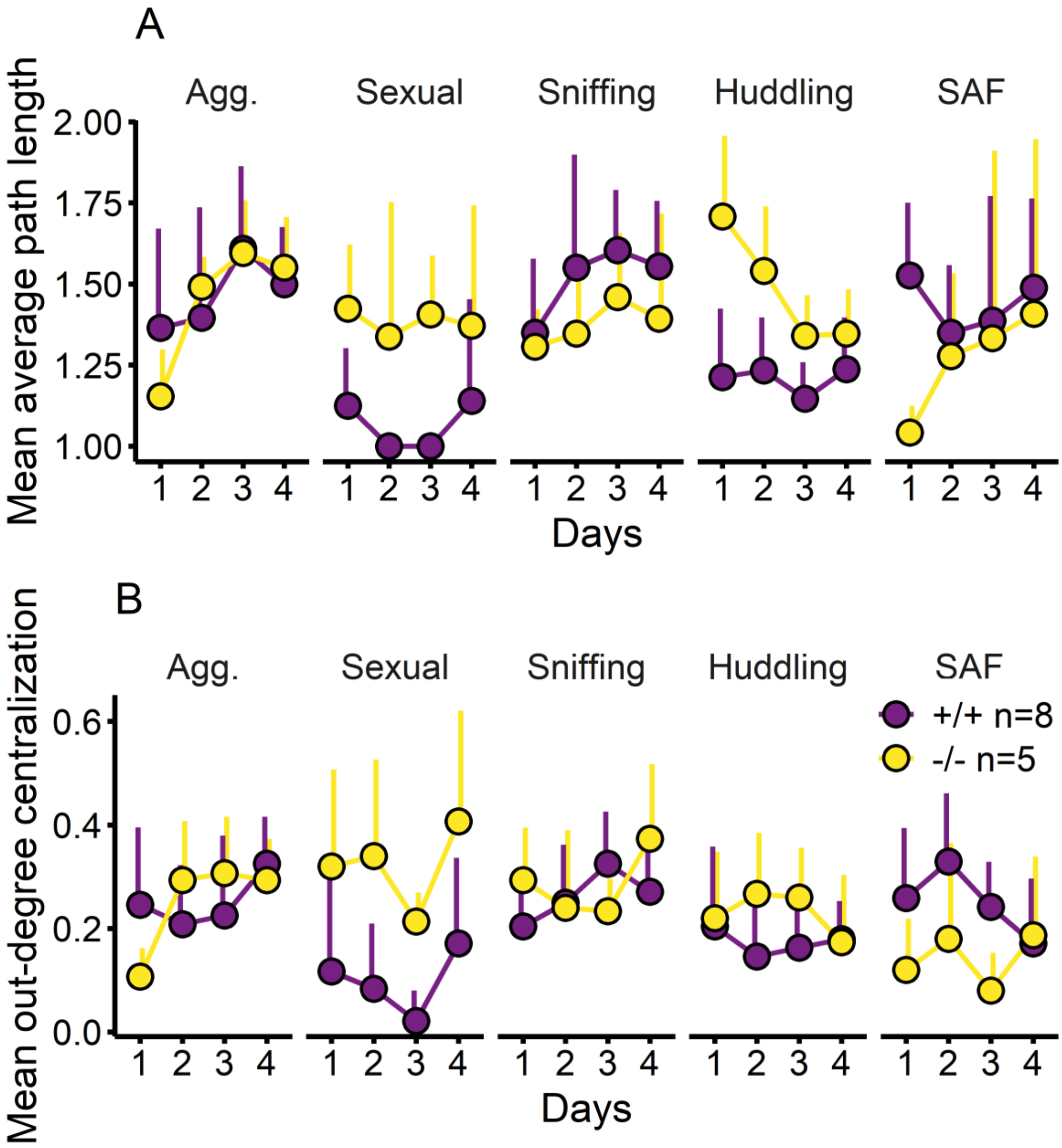

**Figure S6. Average path length and out-degree centralization. A-**Mean average path length for each group of behavior. **B-**Mean out-degree centralization for each group of behavior. "Agg." for general aggression, "SAF" for struggling at the feeder. The global parameters are calculated for the VBS groups. Lines indicate mean + SD. *Tph2*$^{+/+}$ in purple (n = 8 groups) and *Tph2*$^{-/-}$ in yellow (n = 5 groups).

#### G. Figure S7: Glicko rating for each VBS group

In all VBS groups hierarchical ranks emerged progressively. All groups but one had one clear dominant animal with a rating higher than 1/3 of the maximum rating contrast of the group after 4 days of VBS experiment. One group had two dominant animals following the same criterion (Fig, S7I).

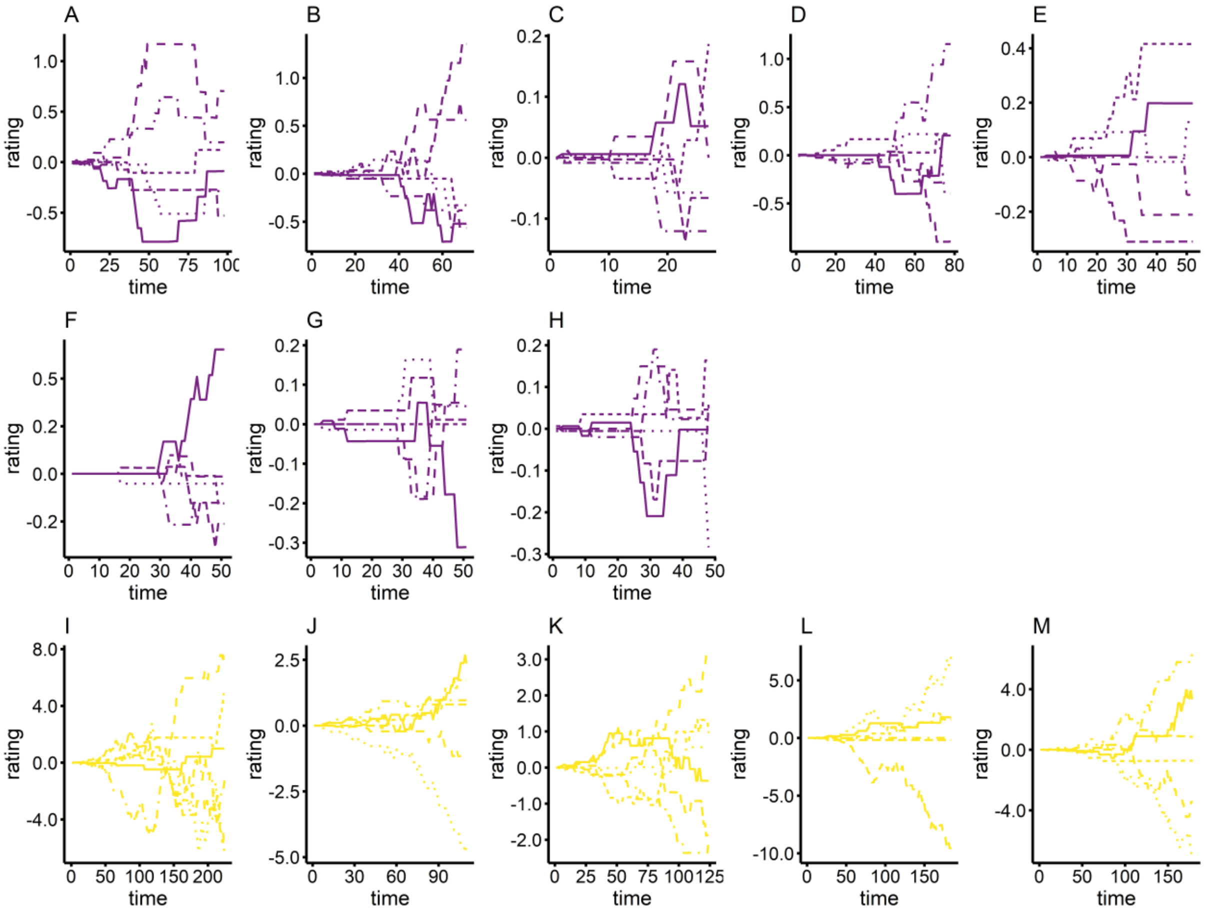

**Figure S7. Glicko rating for each VBS group.** *Tph2*$^{+/+}$ in purple (panels A-H) and *Tph2*$^{-/-}$ in yellow (panels I-M).

### 3. SUPPLEMENTAL TABLES

#### A. Table S1: Number of individuals in each test

Table S1. Number of individuals used for each tests and comment on the exclusion.

| **Test** | ***Tph2*** $^{+/+}$ | ***Tph2*** $^{-/-}$ | **Comment** |
|---|---|---|---|
| Total number | n = 48 | n = 30 | 8 cohorts and 5 cohorts of 6 animals each. |
| RGT | n = 47 | n = 30 | One *Tph2* $^{+/+}$ animal was excluded from the Rat Gambling Task because it did not sample the options. |
| Reversed-RGT | n = 47 | n = 30 | One *Tph2* $^{+/+}$ animal was excluded from the Rat Gambling Task because it did not sample the options; its score in the Reversed-RGT was excluded too. |
| DL-box | n = 24 | n = 24 | Only 4 *Tph2* $^{+/+}$ and 4 *Tph2* $^{-/-}$ cohorts were used. |
| Feces collection | n = 48 | n = 30 | |
| VBS | n = 48 | n = 30 | |
| DDT | n = 48 | n = 30 | |
| SRt | n = 30 | n = 30 | Only 6 *Tph2* $^{+/+}$ and 6 *Tph2* $^{-/-}$ cohorts were used. |
| Odor test | n = 24 | n = 23 | Only 4 *Tph2* $^{+/+}$ and 4 *Tph2* $^{-/-}$ cohorts were used. One *Tph2* $^{-/-}$ animal did not explore the open field (or the odors), its missing value was replaced by the median value of the group. |
| FIEXT | n = 42 | n = 24 | Only 7 *Tph2* $^{+/+}$ and 4 *Tph2* $^{-/-}$ cohorts were used. |
| PDT | n = 24 | n = 24 | Only 4 *Tph2* $^{+/+}$ and 4 *Tph2* $^{-/-}$ cohorts were used. |

#### B. Table S2: One sample t-test for Figure 2A

Table S2. One sample t-test (mean preference compared to 50%) for Figure 2A.

| Genotype | Time point | 0.95 CI | p-value |
|---|---|---|---|
| +/+ | 10 | [43.8, 56.2] | 0.9965 |
| −/− | 10 | [48.8, 67.3] | 0.0853 |
| −/− | 20 | [53.7, 73.9] | 0.008 |
| −/− | 20 | [59.5, 85.2] | < 0.001 |
| +/+ | 30 | [53.6, 75.8] | 0.010 |
| −/− | 30 | [66.2, 90.9] | < 0.001 |
| +/+ | 40 | [60.8, 82.4] | < 0.001 |
| −/− | 40 | [68.4, 91.6] | < 0.001 |
| +/+ | 50 | [62.2, 83.8] | < 0.001 |
| −/− | 50 | [66.1, 91.8] | < 0.001 |
| +/+ | 60 | [59.2, 81.1] | < 0.001 |
| −/− | 60 | [60.4, 87.7] | 0.001 |

#### C. Table S3: Comparison of GDM and PDM in all tests

Table S3. All effect sizes (cohen's d) of GDM vs PDM in *Tph2*$^{+/+}$ and *Tph2*$^{-/-}$ for all other tests than RGT and reversed-RGT.

| Trait | Test | Parameter | *Tph2*$^{+/+}$ GDM vs. PDM cohen's d | *Tph2*$^{-/-}$ GDM vs. PDM cohen's d |
|---|---|---|---|---|
| Impulsive decision making | DDT | AUC DDT | -0.2857334 (small) | -0.3423988 (small) |
| Risky decision making | PDT | AUC PDT | -0.3551802 (small) | -0.07413781 (negligible) |
| Anticipator activity | FI | Mean number of responses | -0.1131997 (negligible) | -0.5166533 (medium) |
| Perseverative activity | EXT | Mean number of responses | -0.3417487 (small) | -0.2303879 (small) |
| Social preference | SRt | Ratio interaction times E1/Hab | 0.6436121 (medium) | 0.1616992 (negligible) |
| Social short term recognition | SRt | Ratio interaction times E1/E3 | 0.3332939 (small) | 0.5761107 (medium) |
| Social long term recognition | SRt | Ratio interaction times Enew/E4 | 0.2659506 (small) | 0.07090615 (negligible) |
| Exploration Dark-Light box | DL box | Time spent in light compartment | -0.3448584 (small) | -0.3876721 (small) |
| Risk taking Dark-Light box | DL box | Risk taking index | 0.7077482 (medium) | 0.6243645 (medium) |
| Activity | VBS | Total distance | 0.005801229 (negligible) | -0.08963622 (negligible) |
| Entropy | VBS | Total roaming entropy | 0.02874787 (negligible) | -0.2498969 (small) |
| Weight loss | VBS | Percentage of weight loss | -0.6171123 (medium) | -0.1698673 (negligible) |
| Corticosterone response | VBS | Percentage of corticosterone after the stay in the VBS | 0.2927969 ( small) | 0.27123 (small) |
| Huddling behavior | VBS | Occurrences huddling behavior | 0.771853 (medium) | -0.218772 (small) |
| Sniffing behavior | VBS | Occurrences sniffing behavior | -0.1976348 (small) | 0.4955432 (small) |
| Eating behavior | VBS | Occurrences eating behavior | -0.1647008 (negligible) | -0.6893535 (medium) |
| Grooming behavior | VBS | Occurrences grooming behavior | 0.04336423 (negligible) | 0.1917227 (negligible) |
| Struggle At Feeder behavior | VBS | Occurrences SAF behavior | 0.1115951 (negligible) | 0 (negligible) |
| General aggression behaviors | VBS | Occurrences gen. aggression behaviors | -0.1386353 (negligible) | -0.4338102 (small) |

| | | | | |
|---|---|---|---|---|
| Sexual behaviors | VBS | Occurrences sexual behaviors | **-0.8021095 (large)** | 0.3501845 (small) |
| Influence over Huddling network | VBS | HUB centrality huddling behavior | **0.8639602 (large)** | 0.1830485 (negligible) |
| Influence over Sniffing network | VBS | HUB centrality sniffing behavior | -0.4372136 (small) | **1.13061 (large)** |
| Influence over SAF network | VBS | HUB centrality SAF behavior | -0.01383219 (negligible) | -0.7042494 (medium) |
| Influence over gen. aggression network | VBS | HUB centrality gen. aggression behaviors | -0.1509906 (negligible) | -0.1867835 (negligible) |
| Influence over Sexual network | VBS | HUB centrality sexual behaviors | -0.5159563 (medium) | -0.02476402 (negligible) |

nota bene: the number of PDM individuals varied from n = 8 to n = 1 in *Tph2*$^{+/+}$ (only n = 3 in the PDT, odor discrimination test and Dark-Light box. and n = 5 in the FIEXT with nose-poke hole and n = 1 in the FIEXT with lever). The number of PDM individuals varied from n = 5 to n = 4 in *Tph2*$^{-/-}$ (one PDM individual excluded in the odor discrimination test).

#### D. Table S4: Wilcoxon rank sum test between for Figure 4A

Table S4. Wilcoxon rank sum test between *Tph2*$^{+/+}$ and *Tph2*$^{-/-}$ for Figure 4A

| **Behavior** | **W** | **p-value** |
|---|---|---|
| Huddling | 1240.5 | **< 0.001** |
| Sniffing | 429 | **0.0028** |
| Eating | 1267 | **< 0.001** |
| Grooming | 914.5 | **0.0459** |
| SAF | 1227.5 | **< 0.001** |
| General aggression | 29 | **< 0.001** |
| Sexual behaviors | 67 | **< 0.001** |

#### E. Table S5: Lmer for Figure 4B

Table S4. Effect of genotype, day and interaction on network density and effect of day on network density for each genotype tested with lmer for Figure 4B

| **Network** | **effect** | **F value (degree of freedom)** | **p-value** |
|---|---|---|---|
| General aggression | genotype | F(1, 43) = 40.9 | **< 0.001** |
| General aggression | day | F(3, 38) = 6.2 | **0.0015** |
| General aggression | genotype x day | F(3, 38) = 7.2 | **< 0.001** |
| General aggression | day (+/+) | F(3, 23) = 1.9 | 0.165 |
| General aggression | day (−/−) | F(3, 12) = 12.6 | **< 0.001** |
| Sexual behaviors | genotype | F(1, 44) = 167 | **< 0.001** |
| Sexual behaviors | day | F(3, 38) = 13.7 | **< 0.001** |
| Sexual behaviors | genotype x day | F(3, 38) = 11 | **< 0.001** |
| Sexual behaviors | day (+/+) | F(3, 23) = 2 | 0.1439 |
| Sexual behaviors | day (−/−) | F(3, 13) = 13 | **< 0.001** |
| Sniffing | genotype | F(1, 32) = 15 | **< 0.001** |
| Sniffing | day | F(3, 22) = 4.7 | **0.0098** |
| Sniffing | genotype x day | F(3, 22) = 1.3 | 0.2943 |
| Sniffing | day (+/+) | F(3, 20) = 9 | **< 0.001** |
| Sniffing | day (−/−) | F(3, 11) = 4 | **0.0365** |
| Huddling | genotype | F(1, 43) = 32.5 | **< 0.001** |
| Huddling | day | F(3, 38) = 5.9 | **0.0019** |
| Huddling | genotype x day | F(3, 38) = 4.7 | **0.0064** |
| Huddling | day (+/+) | F(3, 23) = 0.2 | 0.8919 |
| Huddling | day (−/−) | F(3, 12) = 8.5 | **0.0027** |
| Struggling at feeder | genotype | F(1, 43) = 15.2 | **< 0.001** |
| Struggling at feeder | day | F(3, 38) = 1.2 | 0.2969 |
| Struggling at feeder | genotype x day | F(3, 38) = 0.8 | 0.4805 |
| Struggling at feeder | day (+/+) | F(3, 23) = 2 | 0.1383 |
| Struggling at feeder | day (−/−) | F(3, 14) = 0.2 | 0.8916 |

##### F. Table S6: Random forests on the three datasets

Table S6. Parameters of the Random forest for the three versions of the datasets.

| Versions of the dataset | Number of *Tph2*$^{+/+}$ | Number of *Tph2*$^{-/-}$ | Number of variables | Explanation | Mean accuracy | SD |
|---|---|---|---|---|---|---|
| 1 | n = 48 | n = 30 | 17 | All individuals, omitting PDT, SRt, DL-box and FI-EXT | 98.53% | 0.54 |
| 2 | n = 23 | n = 24 | 24 | omitting FI-EXT | 100% | 0 |
| 3 | n = 17 | n = 18 | 26 | All variables | 100% | 0 |

##### G. Table S7: Principal Component Analysis on the three datasets

Table S7. Parameters of the Principal Component Analysis for the three versions of the datasets. PC for principal component (or dimension).

| Versions of the dataset | Number of *Tph2*$^{+/+}$ | Number of *Tph2*$^{-/-}$ | Number of variables | Explanation | Variance for PC1 (%) | Number of PC to reach 80% |
|---|---|---|---|---|---|---|
| 1 | n = 48 | n = 30 | 17 | All individuals, omitting PDT, SRt, DL-box and FIEXT | 23.13% | 8 PC |
| 2 | n = 23 | n = 24 | 24 | omitting FIEXT | 21.28% | 10 PC |
| 3 | n = 17 | n = 18 | 26 | All variables | 18.71% | 10 PC |

#### H. Table S8: Gini indexes of the Random forests

Table S7. Order of variables per importance (Mean Gini index) for the three versions of the datasets. Most important variables in bold. Variables from classical tests (not VBS) in grey. Total occurrences of sexual behaviors (Sexual), percentage of weight variation (Weight), percentage of corticosterone metabolites variation (Corticosterone), total distance traveled (Distance), total occurrences of defensive behaviors (Defensive), total roaming entropy (Entropy), total occurrences of maintenance behaviors (Maintenance), total occurrences of aggressive behaviors (Aggressive), total preference for open area (Pref.open area), (Affiliative) total occurrences of affiliative behaviors, (AUC.DDT) area under the curve in the DDT, Hub centrality in aggression network (HUB.agg.), flexibility score in reversed-RGT (Flexibility), preference in last 20 min of RGT (RGT), latency to collect pellet in RGT (Latency RGT), Blanchard dominance score (Blanchard), time spent in the light compartment of the DL-box (timeL.DL), index of risk taking in the DL-box test (Risk.taking.DL), area under the curve in the PDT (AUC.PDT), social preference ratio (Socpref), social preference ratio on day 2 of SRt (Socpref.day2), short-term social recognition memory (STM), long-term social recognition memory (LTM), total number of responses in the fixed-interval of FIEXT (FI), total number of responses in the extinction of FIEXT (EXT).

| Version 1 | | | Version 2 | | | Version 3 | | |
|---|---|---|---|---|---|---|---|---|
| | Variable | Mean Gini | | Variable | Mean Gini | | Variable | Mean Gini |
| 1 | **Sexual** | **7.8004641** | 1 | **Corticosterone** | **3.82187434** | 1 | **Corticosterone** | **3.61567559** |
| 2 | **Weight** | **7.7982767** | 2 | Glicko rating | 3.56926427 | 2 | **Affiliative** | **2.97762276** |
| 3 | Corticosterone | 3.9158579 | 3 | Affiliative | 3.44096951 | 3 | Glicko rating | 2.14805505 |
| 4 | Distance | 3.3537214 | 4 | Blanchard | 2.41522225 | 4 | Pref.open area | 2.04673183 |
| 5 | Entropy | 2.8193245 | 5 | Pref.open area | 1.8968134 | 5 | Blanchard | 1.37556659 |
| 6 | Defensive | 2.6518695 | 6 | Sexual | 1.72011537 | 6 | Sexual | 0.76658915 |
| 7 | Maintenance | 2.0825444 | 7 | Defensive | 0.9881357 | 7 | Defensive | 0.60835491 |
| 8 | Glicko rating | 1.4775315 | 8 | Distance | 0.96160999 | 8 | Entropy | 0.51658714 |
| 9 | Aggressive | 1.242993 | 9 | Entropy | 0.90738647 | 9 | HUB.agg. | 0.47006703 |
| 10 | Pref.open area | 1.0495323 | 10 | HUB.agg. | 0.76355717 | 10 | Distance | 0.31380522 |
| 11 | Affiliative | 0.879558 | 11 | timeL.DL | 0.31995666 | 11 | Maintenance | 0.2979113 |
| 12 | AUC.DDT | 0.3668323 | 12 | Socpref | 0.26738162 | 12 | AUC.PDT | 0.26733996 |
| 13 | HUB.agg. | 0.2483838 | 13 | Risk.taking.DL | 0.24820501 | 13 | timeL.DL | 0.1686874 |
| 14 | Flexibility | 0.2315216 | 14 | Weight | 0.24772993 | 14 | EXT | 0.1530195 |
| 15 | RGT | 0.2251097 | 15 | Maintenance | 0.24716998 | 15 | Latency RGT | 0.14738444 |
| 16 | Latency RGT | 0.1886158 | 16 | AUC.PDT | 0.20668295 | 16 | FI | 0.14641028 |
| 17 | Blanchard | 0.1310371 | 17 | LTM | 0.16000447 | 17 | Risk.taking.DL | 0.14597387 |
| | | | 18 | Latency RGT | 0.15219654 | 18 | RGT | 0.12270368 |
| | | | 19 | Flexibility | 0.12945049 | 19 | Socpref.day2 | 0.09864794 |
| | | | 20 | Socpref.day2 | 0.12204077 | 20 | Aggressive | 0.09359775 |
| | | | 21 | AUC.DDT | 0.1074535 | 21 | Weight | 0.09181905 |

| | | | | | | | | |
|---|---|---|---|---|---|---|---|---|
| | | | 22 | STM | 0.10264048 | 22 | LTM | 0.09012947 |
| | | | 23 | Aggressive | 0.09920503 | 23 | Flexibility | 0.08875745 |
| | | | 24 | RGT | 0.09322517 | 24 | STM | 0.08836513 |
| | | | | | | 25 | AUC.DDT | 0.07651249 |
| | | | | | | 26 | Socpref | 0.06485988 |

#### I. Table S9: Contribution of the variables to PC1.

Table S9. Order of variables per contribution to Dimension 1 (PC1) of the Principal Component Analysis (rotation) for the three versions of the datasets. Most important variables in bold. Variables from classical tests in grey. See abbreviations in the text of Table S8.

| Version 1 | | | Version 2 | | | Version 3 | | |
|---|---|---|---|---|---|---|---|---|
| | Variables | Rotation PC1 | | Variables | Rotation PC1 | | Variables | Rotation PC1 |
| 1 | **Weight** | **-0.436** | 1 | **Sexual** | **-0.351** | 1 | **Corticosterone** | **0.358** |
| 2 | **Maintenance** | **0.362** | 2 | **Affiliative** | **-0.344** | 2 | **Pref.open area** | **0.349** |
| 3 | **Entropy** | **0.353** | 3 | **Corticosterone** | **-0.318** | 3 | **Affiliative** | **0.336** |
| 4 | **Corticosterone** | **-0.348** | 4 | **timeL.DL** | **-0.308** | 4 | **Sexual** | **0.315** |
| 5 | **Defensive** | **-0.327** | 5 | Distance | -0.290 | 5 | Defensive | -0.257 |
| 6 | **Sexual** | **-0.325** | 6 | Socpref | -0.283 | 6 | Glicko rating | -0.254 |
| 7 | Distance | -0.259 | 7 | Blanchard | -0.265 | 7 | timeL.DL | 0.254 |
| 8 | Aggressive | -0.251 | 8 | Pref.open area | 0.264 | 8 | Distance | 0.254 |
| 9 | Pref.open area | 0.146 | 9 | Risk.taking.DL | 0.237 | 9 | FI | 0.245 |
| 10 | Flexibility | -0.127 | 10 | Defensive | 0.228 | 10 | Blanchard | 0.242 |
| 11 | Blanchard | 0.124 | 11 | Glicko rating | 0.211 | 11 | Risk.taking.DL | -0.218 |
| 12 | RGT | -0.113 | 12 | Weight | -0.194 | 12 | AUC.PDT | -0.184 |
| 13 | Affiliative | 0.105 | 13 | LTM | -0.157 | 13 | EXT | -0.138 |
| 14 | AUC.DDT | 0.089 | 14 | STM | -0.109 | 14 | Socpref | -0.119 |
| 15 | HUB.agg. | -0.054 | 15 | AUC.PDT | 0.089 | 15 | LTM | 0.105 |
| 16 | Latency RGT | -0.052 | 16 | Latency RGT | 0.082 | 16 | Entropy | -0.096 |
| 17 | Glicko rating | -0.021 | 17 | Entropy | 0.082 | 17 | Weight | 0.094 |
| | | | 18 | Aggressive | 0.077 | 18 | RGT | -0.080 |
| | | | 19 | Socpref.day2 | -0.051 | 19 | Latency RGT | -0.068 |
| | | | 20 | HUB.agg. | -0.025 | 20 | Flexibility | 0.045 |
| | | | 21 | AUC.DDT | 0.012 | 21 | HUB.agg. | 0.041 |
| | | | 22 | Maintenance | -0.005 | 22 | Socpref.day2 | 0.030 |
| | | | 23 | Flexibility | 0.003 | 23 | Aggressive | 0.027 |
| | | | 24 | RGT | 0.003 | 24 | AUC.DDT | -0.014 |
| | | | | | | 25 | STM | -0.010 |
| | | | | | | 26 | Maintenance | -0.005 |

#### J. Table S10: Comparison between VBS impairments and descriptions of human symptoms of mental disorders.

Table S10. Comparison between VBS impairments and descriptions of human symptoms of mental disorders. Impulse control disorders (ICD) are associated with anxiety disorders, obsessive-compulsive disorder (OCD), depression, ADHD, Tourette syndrome and Parkinson's disease (28–33). The first 4 VBS parameters resemble ICDs in human. The last two parameters are more indicative of human stress and anxiety disorders which share a high level of comorbidity with ICDs. See abbreviations in the text of Table S8.

| VBS parameter | Corresponding human parameter | Disease |
|---|---|---|
| **Sexual behavior** | Uncontrollable repetitive sexual behavior, repetition of aggression | ICD (34,35) |
| **Maintenance behavior** | Neglect of personal care<br>(due to repetitive behavior) | ICD (34,35) |
| **Weight** | Neglect of health and personal care (due to repetitive behavior) | ICD (34,35) |
| **Corticosterone** | Cortisol disturbances<br>Stress triggering impulses | ICD (30,36–40)<br>Alcohol use disorder, Borderline personality disorder |
| **Entropy**<br>(as part of hypervigilance profile including smaller territory (roaming entropy) excluding food zones (place preference), higher activity (distance), inhibition maintenance behaviors and huddling, increased stress (corticosterone) | Hypervigilance, attentional bias of vigilance (faster focus of attention on threatening stimuli or persistent focus of attention on threatening stimuli), repetitive checking behavior, generalization of fear responses (over-reaction to harmless stimuli disturbing daily life, avoidance of situations with negative expectation) | OCD, Post-traumatic stress disorder (PTSD), generalized anxiety disorder (28,31,41,42)<br>ICD (43) |
| **Defensive behavior** | Hypervigilance, attentional bias of vigilance (faster focus of attention on threatening stimuli or persistent focus of attention on threatening stimuli), repetitive checking behavior, generalization of fear responses (over-reaction to harmless stimuli disturbing daily life, avoidance of situations with negative expectation) | OCD, PTSD, generalized anxiety disorder (28,31,41,42)<br>ICD (43) |
